# FtsN activates septal cell wall synthesis by forming a processive complex with the septum-specific peptidoglycan synthase in *E. coli*

**DOI:** 10.1101/2021.08.23.457437

**Authors:** Zhixin Lyu, Atsushi Yahashiri, Xinxing Yang, Joshua W. McCausland, Gabriela M. Kaus, Ryan McQuillen, David S. Weiss, Jie Xiao

**Affiliations:** Department of Biophysics and Biophysical Chemistry, Johns Hopkins School of Medicine, Baltimore, MD 21205, USA; Department of Microbiology and Immunology, University of Iowa Carver College of Medicine, Iowa City, IA 52242, USA; Hefei National Laboratory for Physical Science at the Microscale, CAS key Laboratory of Innate Immunity and Chronic Disease, School of Basic Medical Sciences, Division of Life Science and Medicine, University of Science and Technology of China, Hefei, China

## Abstract

The FtsN protein of *Escherichia coli* and other proteobacteria is an essential and highly conserved bitopic membrane protein that triggers the inward synthesis of septal peptidoglycan (sPG) during cell division. Previous work has shown that the activation of sPG synthesis by FtsN involves a series of interactions of FtsN with other divisome proteins and the cell wall. Precisely how FtsN achieves this role is unclear, but a recent study has shown that FtsN promotes the relocation of the essential sPG synthase FtsWI from an FtsZ-associated track (where FtsWI is inactive) to an sPG-track (where FtsWI engages in sPG synthesis). Whether FtsN works by displacing FtsWI from the Z-track or capturing/retaining FtsWI on the sPG-track is not known. Here we use single-molecule imaging and genetic manipulation to investigate the organization and dynamics of FtsN at the septum and how they are coupled to sPG synthesis activity. We found that FtsN exhibits a spatial organization and dynamics distinct from those of the FtsZ-ring. Single FtsN molecules move processively as a single population with a speed of ∼ 9 nm s^-1^, similar to the speed of active FtsWI molecules on the sPG-track, but significantly different from the ∼ 30 nm s^-1^ speed of inactive FtsWI molecules on the FtsZ-track. Furthermore, the processive movement of FtsN is independent of FtsZ’s treadmilling dynamics but driven exclusively by active sPG synthesis. Importantly, only the essential domain of FtsN, a three-helix bundle in the periplasm, is required to maintain the processive complex containing both FtsWI and FtsN on the sPG-track. We conclude that FtsN activates sPG synthesis by forming a processive synthesis complex with FtsWI exclusively on the sPG-track. These findings favor a model in which FtsN captures or retains FtsWI on the sPG-track rather than one in which FtsN actively displaces FtsWI from the Z-track.

## Introduction

Most bacteria are completely encased in a peptidoglycan (PG) sacculus (or cell wall). The cell wall confers cell shape and protects against lysis by high internal osmotic pressure (up to ∼ 3 atm in Gram-negative *Escherichia coli*^1^ and 20 atm in Gram-positive *Bacillus subtilis*^2^). The importance of the cell wall is underscored by the fact that it is one of the most successful antibiotic targets^3, 4^.

During cell division bacteria must synthesize and remodel their protective cell wall to accommodate the splitting of a mother cell into two daughter cells^5^. Bacterial cell division is mediated by the divisome, a loosely-defined collection of proteins that form a contractile ring-like assemblage at the division site. In *E. coli* the divisome contains over 30 different types of proteins, of which ten are essential and considered to constitute the core of the division apparatus^6, 7^. The ten essential division proteins are recruited to the divisome in a mostly sequential fashion, starting with the tubulin-like GTPase FtsZ^8, 9^ and ending with the bitopic membrane protein FtsN, whose arrival coincides with the onset of visible septum constriction^10, 11^. Other noteworthy divisome proteins include FtsA, which links FtsZ polymers to the membrane^12^, the core septal PG (sPG) synthase complex composed of the polymerase FtsW and transpeptidase FtsI^13, 14^, and the FtsQLB complex, which regulates FtsWI activity^15, 16^ (also see reviews^5-7^). According to current models, FtsN acts through FtsA in the cytoplasm and the FtsQLB complex in the periplasm to activate synthesis of sPG by the FtsWI synthase complex^15-21^.

Although a small amount of FtsN is recruited early during divisome assembly by binding directly to FtsA, the majority of septal FtsN localizes later *via* binding of its C-terminal SPOR domain to the glycan backbone of PG at sites that lack stem peptides^11, 17, 22-24^. The SPOR domain binding sites, referred to here as “denuded” glycans, are a hallmark of sPG. Denuded glycans are created by cell wall amidases that process sPG to allow for daughter cell separation^25^ and are subsequently degraded by lytic transglycosylases^26^ (also see reviews^5, 27, 28^). Thus, denuded glycans accumulate only transiently in sPG.

Advanced high resolution and single-molecule imaging are providing important new insights into the organization of the divisome and the control of sPG synthesis^29-36^. One of these recent insights is that FtsZ uses GTP hydrolysis to move around the septum by treadmilling^37-40^, which is the apparent directional movement of a polymer caused by the continuous polymerization at one end and depolymerization at the other end, with individual monomers in the middle remaining stationary. Furthermore, in both *E. coli* and *B. subtilis*, FtsZ’s treadmilling dynamics were found to drive the directional movement of the sPG synthesis complex FtsWI at a speed of ∼ 30 nm s^-1^ ^37, 38^, likely through a Brownian ratchet mechanism^41^. Thus, FtsZ uses its GTPase activity-dependent treadmilling dynamics to function as a linear motor to distribute sPG enzyme complexes along the septum to ensure a smooth, symmetric septum synthesis^29, 37, 38, 41^.

More recently, we discovered that the *E. coli* divisome contains a second population of FtsWI, one that moves processively at ∼ 8-9 nm s^-1^ ^42^. Movement of this slower population is driven by active PG synthesis (termed as on the sPG-track) rather than FtsZ treadmilling (termed as on the Z-track). Similar FtsZ-independent but sPG synthesis-dependent processive populations of FtsW and PBP2x were first observed in *S. pneumoniae*^40^. In *E. coli* individual FtsW or FtsI molecules can transition back-and-forth between the fast- and slow-moving populations. In cells depleted of FtsN, the active, slow-moving population of FtsWI on the sPG-track is diminished while the inactive, fast-moving population of FtsWI on the Z-track is enhanced. These findings imply that FtsN activates sPG synthesis, at least in part, by increasing the number of FtsWI molecules on the sPG-track^42^. Whether FtsN accomplishes this by releasing FtsWI from the Z-track or retaining FtsWI on the sPG-track is not yet known.

In this work, we use single-molecule imaging to investigate the organization and dynamics of FtsN at the septum and how they are coupled to sPG synthesis. We found that FtsN exhibits distinct spatial organization and dynamics from the FtsZ-ring, supporting the previous notion that the FtsN-ring organization is independent of the FtsZ-ring^43^. Most importantly, we observed that single FtsN molecules move exclusively and processively at a slow speed of ∼ 9 nm s^-1^ at the septum. The processive movement of FtsN depends on active sPG synthesis but not FtsZ’s treadmilling dynamics. These dynamic behaviors are identical to those of the slow-moving, active population of FtsWI. Moreover, only the essential domain of FtsN, a helix bundle in the periplasm, is required for processive movement of FtsN on the sPG-track. These findings support a model whereby FtsN activates sPG synthesis by forming a processive complex with FtsWI through the essential domain to maintain it on the sPG-track and thus promote the synthesis of sPG.

## Results

### Construction of functional FtsN fusions

FtsN has at least four functional domains (**Figure 1A, Figure S1**): an N-terminal cytoplasmic tail (FtsN^Cyto^) that interacts with FtsA^17, 19, 20^; a transmembrane domain (FtsN™) that anchors FtsN to the inner membrane^44^; a periplasmic essential domain (FtsN^E^) that is composed of three helices and responsible for activating sPG synthesis activity^11, 45^, and a C-terminal periplasmic SPOR domain (FtsN^SPOR^) that binds to denuded glycan strands, which are transiently present at the septum during cell wall constriction^11, 22^-^24, 45, 46^. To identify functional fluorescent fusions of FtsN, we designed and screened 11 FtsN fusions that have a green fluorescent protein mNeonGreen (mNeG)^47^ fused to the N-terminus, C-terminus or inserted at internal positions of FtsN (**Figure S1**). These fusions were expressed from plasmids in an FtsN-depletion background to test their functionality (**Figure S2**). We were able to identify an N-terminal and an internal (termed sandwich, between E60 and E61) fusion of FtsN that supported normal growth on solid and liquid media in FtsN depletion backgrounds (**Figure S2A, B**), and exhibited correct midcell localization during cell division (**Figure S2C**). Based on these results we constructed additional fusions to various fluorescent proteins for different imaging purposes, including the N-terminal fusions mEos3.2-FtsN, GFP-FtsN, and the sandwich fusion FtsN-Halo^SW^ (**Figure S3** and **Supplemental Notes**). These fusions were integrated into the chromosome at a phage attachment site in an FtsN-depletion strain constructed by replacing *ftsN’s* native promoter with the arabinose-dependent *P*_*BAD*_ promoter^48^ (**Table S1**). Expression, stability and functionality of these fusions were further validated by Western blotting and cell growth measurements (**Figure S3**).

**Figure 1.**
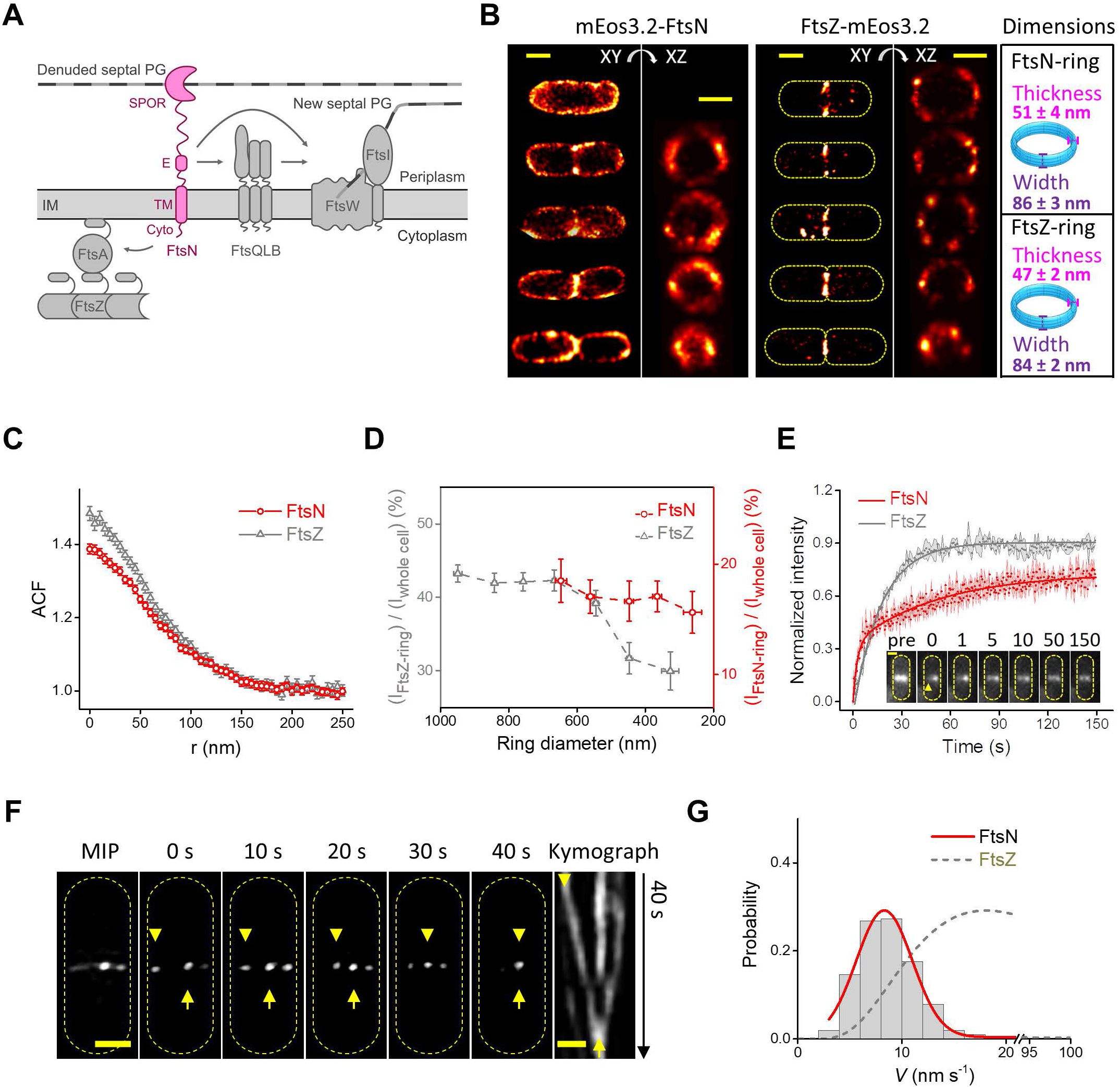
FtsN-ring has different organization and dynamics compared to FtsZ-ring. (A) Schematic drawing of FtsN’s domain organization and interaction with other divisome proteins. FtsN is recruited to and localizes to the division site through the interaction of its cytoplasmic domain (Cyto) with FtsA and its periplasmic SPOR domain with septal peptidoglycan (sPG). The sPG synthase complex FtsWI is activated by FtsN’s essential domain (E) either directly or indirectly via FtsQLB. (B) Three-dimensional (3D) live cell single-molecule localization superresolution microscopy (SMLM) images show FtsN-rings (left) are patchy but more homogenously than FtsZ-rings (middle). Rings with similar diameters and localization numbers are chosen for comparison. Yellow dashes mark cell outlines for illustrative purpose. Scale bars, 500 nm. Toroid ring models (cyan) are shown on the right, with the average ring width along the long axis of cells at 86 ± 3 nm and average radial thickness at 51 ± 4 nm, *n* = 72 rings for FtsN-rings, and width at 84 ± 2 nm and radial thickness at 47 ± 2 nm, *n* = 103 rings for FtsZ-rings. All measurements are expressed as mean ± standard error of the mean, *μ* ± *s*.*e*.*m*. (C) Mean spatial autocorrelation function (ACF) curves of FtsN-rings (red open circle, *n* = 72 cells) and FtsZ-rings (gray open triangle, *n* = 103 cells) averaged from all individual cells’ ACFs. x-axis (r) is the distance between each molecule pair. Error bars represent *s*.*e*.*m*. The lower correlation *v*alue at short distances and longer characteristic decay length of FtsN’s ACF indicate a more homogenous distribution of FtsN in the rings compared to that of FtsZ. (D) A pseudo time course of FtsN’s (red open circle) and FtsZ’s (gray open triangle) midcell localization percentages (*I*_ring_/*I*_whole cell_) during cell division. FtsN assemble and disassemble at a later stage of cell division than FtsZ. (E) Mean FRAP recovery curve of FtsN (red, *n* = 58 cells) exhibits slower and lower recovery than that of FtsZ (gray, data from a previous work^38^). Examples of raw FRAP images of a GFP-FtsN expressing cell (Strain EC4240 in **Table S1**) are shown as inset (arrowhead shows the bleaching area, **Figure S6A**). Scale bar, 300 nm. (F) Maximum intensity projection (MIP, left) and montages (0-40 s) from time-lapse imaging of a cell with a clearly visible midcell FtsN-ring using TIRF-SIM. Scale bar, 300 nm. Kymograph is compiled from the fluorescent intensity along the septum over time. The arrowhead points to a moving cluster while the arrow points to a stationary cluster. Scale bar, 200 nm. (G) Speed distribution of processively moving FtsN clusters combined from both TIRF-SIM and TIRF imaging (gray columns) overlaid with the corresponding fit curve (solid red) and a fit curve of FtsZ’s treadmilling speed distribution (dash gray, data from a previous work^42^). The average moving speed of combined FtsN clusters is 8.7 ± 0.2 nm s^-1^, *μ* ± *s*.*e*.*m*., *n* = 205 clusters. A break of the *x*-axis from 21 to 94 nm s^-1^ was used to accommodate the distinct speed distributions between FtsN and FtsZ clusters.

### The FtsN-ring exhibits different spatiotemporal organization and dynamics from the FtsZ-ring

FtsN is expressed at a level of ∼ 300 molecules per cell^49^ (**Figure S4**) and forms a ring-like structure (FtsN-ring) at the midcell similar to the FtsZ-ring^43^. To understand the spatial organization of the FtsN-ring, we performed astigmatism-based three-dimensional (3D) single-molecule localization microscopy (SMLM)^50^ imaging in live *E. coli* cells using an mEos3.2-FtsN fusion (Strain EC4443 in **Table S1**). Under our imaging condition, we achieved a spatial resolution of ∼ 50 nm in the *xy* axis and ∼ 80 nm in *z* axis (**Figure S5B**). Similar to what we and others have observed for the FtsZ-ring^33, 35, 36, 51, 52^, FtsN-rings are patchy and have comparable dimensions to FtsZ-rings (**Figure 1B, Figure S5C**, and **Table S5**). However, autocorrelation analysis showed that the FtsN molecules in the FtsN-ring are more homogenously distributed than those in the FtsZ-ring (**Figure 1C**), indicating a different spatial organization of the FtsN-ring.

Next, we sorted individual cells by their ring diameters to generate a pseudo time lapse representing the cell wall constriction process. We found that FtsN-rings assemble at a ring diameter of ∼ 600 nm and disassemble at ∼ 300 nm (**Figure 1D**). In contrast, under the same experimental condition FtsZ-rings assemble at ∼ 950 nm and start to disassemble at ∼ 600 nm (**Figure 1D**). These results demonstrate that the FtsN-ring assembles and disassembles at cell wall constriction stages significantly later than the Z-ring.

We next investigated whether the FtsN-ring exhibits similar dynamic subunit turnovers as were observed for the FtsZ-ring^53, 54^. To do so we carried out Fluorescence Recovery After Photobleaching (FRAP) experiments using a GFP-FtsN fusion (**Figure S6, Supplemental Movie 1**, Strain EC4240 in **Table S1**). By bleaching half of the ring, we found that the recovery curve of GFP-FtsN exhibited two apparent phases (**Figure 1E**), a fast phase with a recovery half time *τ*_1/2_ = 2.9 ± 0.8 s, and a slow phase with *τ*_1/2_ = 54 ± 10 s (*μ* ± *s*.*e*.*m*., *n* = 58 cells). Most interestingly, we only observed a ∼ 70% recovery of FtsN’s intensity compared to that prior to bleaching, indicating that a population of FtsN molecules were stationary on the time scale of the experiment (150 s). In comparison, at the same time scale the FtsZ-ring recovered with a half time of ∼ 16 s and to ∼ 90% of the intensity prior to bleaching (**Figure 1E**, data from a previous work^38^). The fast recovery phase of FtsN was also previously observed by Söderström *et al*.^43^, which is most likely due to the random diffusion of FtsN molecules in and out of the septum as expected for a typical inner membrane protein (**Supplemental Notes**). The slow recovery phase, however, is significantly slower than that of FtsZ, indicating that FtsN-ring exhibits different dynamics compared to the FtsZ-ring.

Taken together, these results are consistent with previous observations that FtsN and FtsZ do not colocalize with each other at the molecular scale revealed by superresolution imaging^43^. They suggest that the spatiotemporal organization and dynamics of FtsN are most likely independent of FtsZ.

### FtsN clusters exhibit slow, directional motions

To investigate what type of dynamics contribute to the observed slow FRAP behavior, we imaged FtsN-rings using an mNeG-FtsN fusion (Strain EC4564 in **Table S1**) with structured illumination microscopy coupled with total internal reflection excitation (TIRF-SIM)^55, 56^. TIRF-SIM allowed us to monitor the dynamics of FtsN-rings with a spatial resolution of ∼ 100 nm and a time resolution of 100 ms. Similar to what we observed in 3D-SMLM imaging, fluorescence of FtsN-rings was patchy and clustered (**Figure 1F, Figure S7B**). Kymograph analysis showed that some FtsN clusters are stationary and remained at the same position throughout the imaging time (40 s, **Figure 1F**, arrow, **Supplemental Movie 2**). These stationary FtsN clusters likely explain the fraction of unrecovered FRAP signal. However, some FtsN clusters exhibited apparently transverse, processive movement across the short axis of the cell (**Figure 1F**, arrowhead, **Supplemental Movie 2**). The mean directional speed measured from these kymographs was at 8.8 ± 0.3 nm s^-1^, (*μ* ± *s*.*e*.*m*., *n* = 92 clusters). These directionally moving FtsN clusters are likely the ones contributing to the slow recovery rate of FRAP, as it takes ∼ 60 s for an FtsN cluster at this speed to cross the TIRF-SIM imaging field (∼ 500 nm, **Supplemental Notes**). We further confirmed that the directional motion was not due to SIM imaging artifacts as we obtained the same result (*v* = 8.6 ± 0.3 nm s^-1^, *μ* ± *s*.*e*.*m*., *n* = 113 clusters) using the same mNeG-FtsN fusion in conventional TIRF imaging even though the spatial resolution was lower (**Figure S7A**). The directional motion was not due to stage drifting either, because we observed both stationary and moving clusters in the same cells (**Figure 1F**). Furthermore, in fixed cells, the directional, processive movement of FtsN was completely abolished (**Figure S7B**). The combined ∼ 9 nm s^-1^ directional moving speed of FtsN clusters (**Table S6**) is significantly slower than the treadmilling speed of FtsZ polymers (∼ 30 nm s^-1^)^37, 38^ (**Figure 1G**), again demonstrating that this motion is distinct from the treadmilling dynamics of FtsZ.

### Individual FtsN molecules exhibit slow, directional motions

Apparent directional motion of a protein cluster can arise from the coordinated directional movement of individual protein molecules in the cluster or treadmilling dynamics. The latter has been reported for a few bacterial cytoskeletal proteins^57, 58^, most recently FtsZ^37-40^ and PhuZ^59^. To distinguish between these two possibilities, we used 3D single-molecule tracking (3D-SMT) to investigate the movement of single FtsN molecules.

To facilitate SMT, we used a FtsN-Halo^SW^ fusion (Strain EC5234 in **Table S1**) that can be sparsely labeled with the bright organic dye JF646 added into the growth medium^60^. The Halo tag is inserted after amino acid E60, between the TM and E domains in the periplasm (**Figure S1**). We tracked septum-localized single FtsN-Halo^SW^ molecules using a frame rate of 1 Hz to effectively filter out fast, randomly diffusing molecules along the cylindrical part of the cell body. Using a custom-developed unwrapping algorithm^41, 42^, we decomposed 3D trajectories of individual FtsN molecules obtained from the curved cell surfaces at midcell to one-dimensional (1D) trajectories along the circumference and long axis of the cell respectively as previously described^42^.

We found that some FtsN molecules were confined to small regions at the septum and stayed stationary (**Figure 2A**). Some moved directionally across the cell’s short axis (**Figure 2B**). Some others dynamically transitioned in between different moving speeds and directions (**Figure 2C**). To quantify these behaviors, we used a trajectory segmentation method we previously described^41, 42^ and classified segments as either stationary or moving directionally based on a statistical criterion (**Supplemental Notes**). We found that, on average, ∼ 55% (55.1 ± 1.6%) of the segments were classified as stationary (**Figure 2D**, solid black) with an average dwell time of ∼ 27 s (27.3 ± 1.3 s, *µ* ± *s*.*e*.*m*., *n* = 315 segments, **Table S10**). For the rest of the segments, FtsN molecule engages in directional movement as a single population (**Figure S8**) at the septum with an average run time of ∼ 15 s (14.5 ± 0.7 s, *μ* ± *s*.*e*.*m*., *n* = 256 segments, **Table S10**) and average run speed of 9.4 ± 0.2 nm s^-1^ (*μ* ± *s*.*e*.*m*., **Figure 2D**, solid red, **Table S10**). Notably, with the two-sample Kolmogorov-Smirnov (K-S) test, the speed distribution is essentially the same as what we observed for mNeG-FtsN clusters using TIRF-SIM (**Figure S9**), similar to what we previously measured for the slow-moving population of active FtsW and FtsI engaged on the sPG-track in our recent studies^42^ (average at 9.4 ± 0.3 nm s^-1^, **Figure 2D**, red dash), and has a minimal overlap with FtsZ’s treadmilling speed distribution under the same condition (average at 28.0 ± 1.2 nm s^-1^, **Figure 2D**, gray dash). Thus, FtsN’s directional movement resembles that of the active, slow-moving population of FtsWI on the sPG-track, but not the inactive, fast-moving population of FtsWI on the FtsZ-track.

**Figure 2.**
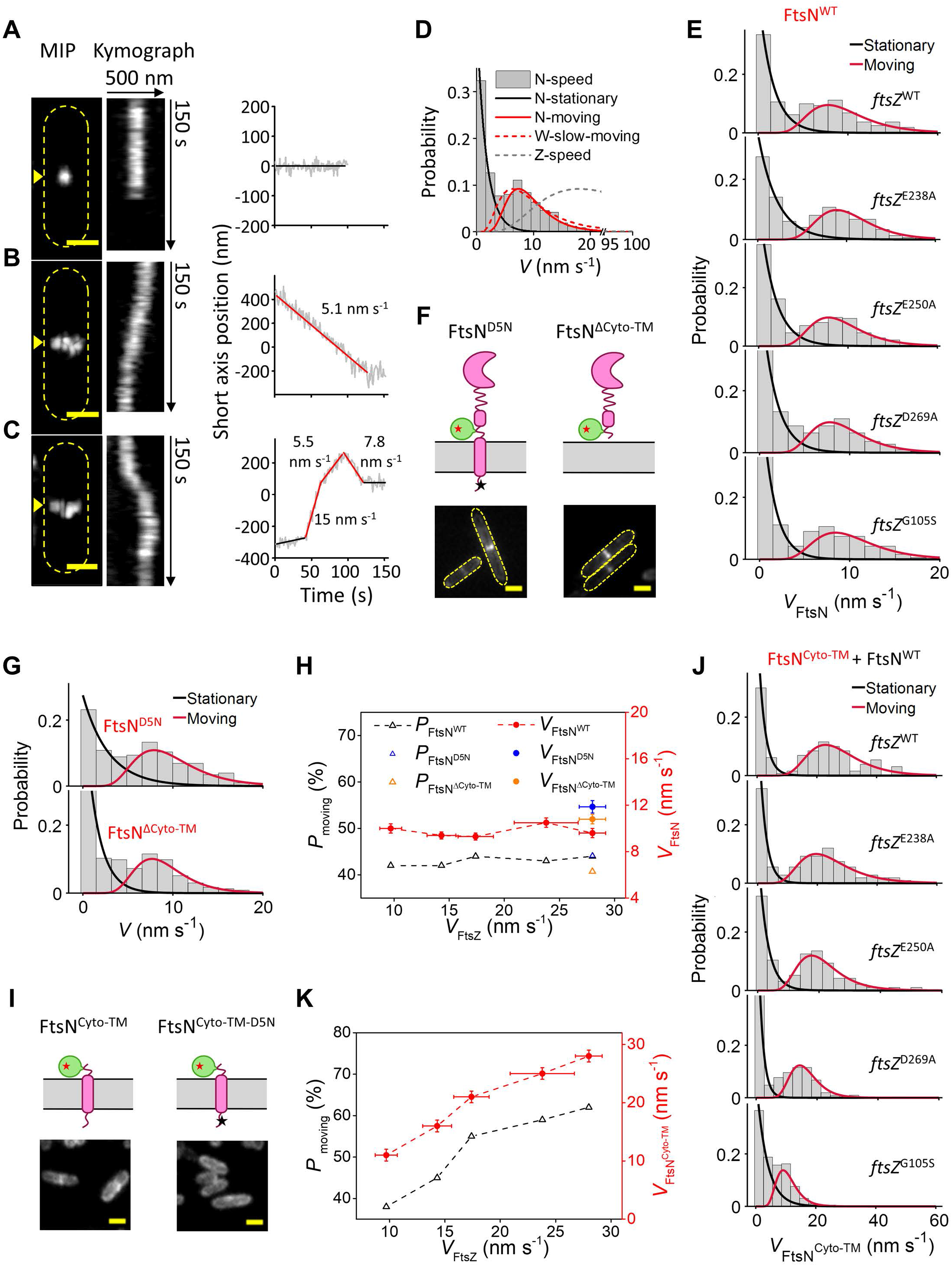
FtsN exhibits a single processive moving population slower than, and independent of, the treadmilling dynamics of FtsZ. (A-C) Representative maximum fluorescence intensity projection images (left), kymographs of fluorescence line scans at the septa (marked by yellow arrow head, middle), and unwrapped one-dimensional positions of the corresponding FtsN-Halo^SW^ molecule (right) along the circumference of the cell of a stationary FtsN-Halo^SW^ molecule (A), a directionally moving FtsN-Halo^SW^ molecule (B), and an FtsN-Halo^SW^ molecule that transitioned between different directions and speeds (C). Each segment was fit by a straight-line and classified as stationary (black) or processively moving (red) based on a statistic criterion. Scale bars, 500 nm. (D) FtsN’s speed distribution (gray columns) overlaid with the fit curves of the stationary (solid black) and moving (solid red) populations. For comparison, the fit curves of the slow-moving population of FtsW molecules (dash red, data from a previous work^42^) and FtsZ’s treadmilling speed distribution (dash gray, data from a previous work^38^) were superimposed. A break of the *x*-axis from 22 to 93 nm s^-1^ was used to accommodate the distinct speed distributions between FtsN and FtsZ. (E) Speed distributions of single FtsN-Halo^SW^ molecules in WT and *ftsZ* GTPase mutant strains overlaid with corresponding fit curves (stationary population in black and moving population in red). (F) Schematic representation of two FtsN mutants, FtsN^D5N^-Halo^SW^ (left) and Halo-FtsN^ΔCyto-TM^ (right). In the FtsN^D5N^-Halo^SW^ mutant, the black star in the cytoplasmic tail represents the D5N mutation. In both mutants, the green bubble represents the Halo tag and the red star represents the JF646 dye. Representative ensemble fluorescence cell images are shown at the bottom. Scale bars, 1 µm. (G) Speed distributions of single FtsN^D5N^-Halo^SW^ (top) and Halo-FtsN^ΔCyto-TM^ (bottom) molecules overlaid with corresponding fit curves. (H) Percentage of moving population (black triangle) and average moving speed (red cycle) of FtsN are independent of FtsZ’s treadmilling speed. FtsN^D5N^ and FtsN^∆Cyto-TM^ data are shown in blue and orange, respectively. (I) Schematic representation of FtsN^Cyto-TM^-Halo^SW^ (left) and FtsN^Cyto-TM-D5N^-Halo^SW^ (right). Representative ensemble fluorescence cell images are shown at the bottom. Scale bars, 1 µm. (J) Speed distributions of single FtsN^Cyto-TM^-Halo^SW^ molecules in WT and *ftsZ* GTPase mutant strains overlaid with corresponding fit curves (stationary population in black and moving population in red). (K) Percentage of moving population (black triangle) and average moving speed (red cycle) of FtsN^Cyto-TM^ are dependent on FtsZ’s treadmilling speed.

### FtsN’s slow, directional movement is independent of FtsZ’s treadmilling dynamics

Our previous studies have shown that the slow-moving population of FtsWI is independent of FtsZ’s treadmilling dynamics but dependent on active sPG synthesis^42^. Because the speed distribution of FtsN largely overlaps with that of the slow-moving population of FtsWI (**Figure 2D**), we reasoned that FtsN likely moves together with FtsWI as part of an active sPG synthesis complex. If so, we would expect that FtsN’s motion depends on active sPG synthesis but not on FtsZ’s treadmilling dynamics in the same manner as FtsWI.

To test whether FtsN’s motion is FtsZ-dependent, we performed SMT of FtsN-Halo^SW^ in four FtsZ GTPase mutant strains which show progressively slower treadmilling speeds (*ftsZ*^*E238A*^, *ftsZ*^*E250A*^, *ftsZ*^*D269A*^, and *ftsZ*^*G105S*^). As expected, the average speed of directionally moving FtsN molecules in these mutants remained constant at ∼ 9 nm s^-1^ (**Figure 2E, Table S7**), independent of FtsZ’s treadmilling speed (**Figure 2E, H**). This behavior is essentially the same as the slow-moving, active population of FtsW and FtsI^42^. Similarly, the percentage of FtsN molecules that were moving directionally remained constant in these mutant backgrounds (**Figure 2E, H**). These results demonstrate that FtsN’s slow-moving dynamics are not driven by FtsZ’s treadmilling dynamics.

### FtsN’s slow, directional movement is independent of its cytoplasmic domain

The independence of FtsN’s directional motion from FtsZ dynamics is somewhat unexpected in light of previous reports that the N-terminal cytoplasmic domain (Cyto) of FtsN can localize to the midcell through its direct interaction with the 1C domain of FtsA^17, 19, 20, 61-65^. To address whether this or any other cytoplasmic interaction contributes to the ∼ 9 nm s^-1^ directional movement of FtsN, we constructed two FtsN mutants (**Figure 2F**). One mutant contains a D5N mutation in the N-terminal cytoplasmic domain (FtsN^D5N^- Halo^SW^, Strain EC5271 in **Table S1**) that has been shown to reduce the interaction between FtsN and FtsA^20^. In the other mutant we deleted the entire cytoplasmic and transmembrane domains, fusing the periplasmic region of FtsN to the cleavable signal sequence from DsbA to export the fusion directly to the periplasm (DsbA^ss^-Halo-FtsN^ΔCyto-TM^, Strain EC5263 in **Table S1**). Both mutants were able to support cell division as the sole cellular FtsN copy expressed from the endogenous chromosomal locus and showed prominent midcell localization, but cells were both longer than WT ones (**Figure 2F, Figure S10**), likely due to delayed initiation or slowed rate of cell wall constriction because of the lack of the N-terminal interactions. Interestingly, both mutants exhibited essentially unchanged percentage or speed of the directionally moving population (**Figure 2G, H, Table S8**). These results strongly suggest that the interactions between the cytoplasmic domain of FtsN and FtsA do not contribute to the observed slow-moving dynamics of FtsN.

### FtsN’s cytoplasmic domain exhibits fast, FtsZ treadmilling-dependent directional movement

Although we did not observe any FtsZ-dependent directional motion of FtsN as what we observed for FtsWI, we reasoned that the cytoplasmic interaction between FtsN and FtsA may still be able to mediate the end-tracking behavior of FtsN on treadmilling FtsZ polymers using a Brownian ratchet mechanism as we previously predicted^41^. This interaction may only exist in cells at an early divisome assembly stage, which were not well represented in the imaging samples, and it may be diminished after FtsN is recruited to the midcell due to the presence of FtsN’s periplasmic interactions with other divisome proteins and/or the denuded glycan strands.

To examine this possibility, we constructed a FtsN^Cyto-TM^-Halo^SW^ fusion, in which the E and SPOR domains of FtsN are removed and a Halo tag is inserted in the same position (E60-E61) as the full-length sandwich fusion (**Figure 2I**, Strain EC5317 in **Table S1**). Because FtsN^Cyto-TM^ cannot support cell division by itself, we expressed it ectopically from the chromosome in the presence of WT FtsN. Ensemble fluorescence imaging showed that FtsN^Cyto-TM^-Halo^SW^ exhibits patchy fluorescence along the cell perimeter and has markedly decreased midcell localization compared to full length FtsN (**Figure 2I**). This observation is consistent with FtsN^Cyto-TM^ having a transmembrane domain but not the SPOR domain, which is the major septum localization determinant^11, 24^. Further mutating the conserved D5 residue in the cytoplasmic domain (FtsN^Cyto-TM-D5N^-Halo^SW^, Strain EC5321 in **Table S1**) completely abolished any residual midcell localization (**Figure 2I**), demonstrating that the limited midcell localization of FtsN^Cyto-TM^-Halo^SW^ is indeed mediated by FtsN’s interaction with FtsA.

Despite the poor septal localization, we were able to track remaining single FtsN^Cyto-TM^- Halo^SW^ molecules at the midcell in a series of FtsZ GTPase WT and mutant backgrounds. Strikingly, we found that now in ∼ 60% (62.5 ± 1.9%) of the SMT segments FtsN^Cyto-TM^- Halo^SW^ molecules moved at an average speed of ∼ 30 nm s^-1^ (29.1 ± 1.7 nm s^-1^, *µ* ± *s*.*e*.*m*., *n* = 130 segments, **Figure 2J, Table S9**) in the FtsZ WT background, similar to FtsZ’s treadmilling speed. In four FtsZ GTPase mutant strains (*ftsZ*^*E238A*^, *ftsZ*^*E250A*^, *ftsZ*^*D269A*^, *and ftsZ*^*G105S*^), we observed progressively reduced speed and population percentage of directionally-moving FtsN^Cyto-TM^-Halo^SW^ (**Figure 2K, Table S9**). There was no discernible slow-moving population of FtsN^Cyto-TM^-Halo^SW^ under any of these conditions. These results strongly suggest that *in vivo* the cytoplasmic interaction between FtsN^Cyto-TM^ and FtsA is able to drive the FtsZ treadmilling-dependent end-tracking behavior of FtsN^Cyto-TM^ and that this interaction is diminished once FtsN’s periplasmic interactions take place during the process of cell division. A previous *in vitro* study showed that membrane-anchored cytoplasmic domain of FtsN is capable of following treadmilling FtsZ polymers through a diffusion-and-capture mechanism^66^, but does not directionally end-track FtsZ at the single-molecule level as what we observed here. This difference is most likely due to the more restricted diffusion of FtsN^Cyto-TM^ along the septum area *in vivo* compared to that *in vitro*, as we previously predicted in a Brownian ratchet model^41^.

### FtsN’s directional movement depends on sPG synthesis

Our results so far demonstrated that the slow, directional movement of full length FtsN is independent of FtsZ’s treadmilling dynamics. To examine whether it is driven by active sPG synthesis as that for the slow-moving population of FtsWI^42^, we performed SMT of FtsN-Halo^SW^ under conditions of altered sPG synthesis activities.

We first examined the effect of inhibiting FtsW’s glycosyltransferase (GTase) activity on the movement of FtsN-Halo^SW^ using a functional FtsW variant, FtsW^I302C^, which can be specifically inhibited upon the addition of the cysteine-reactive reagent MTSES (2-sulfonatoethylmethanethiosulfonate)^42^. In this strain background, FtsN-Halo^SW^ exhibited similar dynamics as in the parent FtsW^WT^ cells (**Figure 3A**, top two panels, **Table S10**). In the presence of MTSES (100 µM, 60 min), however, the directionally moving population of FtsN-Halo^SW^ was significantly reduced and on average ∼ 80% of segments were stationary (80.4 ± 1.4%, *n* = 115 segments, **Figure 3A, C, Table S10**). The depletion of the moving population is essentially identical to the depletion of the slow-moving population of FtsW^I302C^ in the presence of MTSES and suggests that the directional motion of FtsN is coupled to FtsW and its GTase activity.

**Figure 3.**
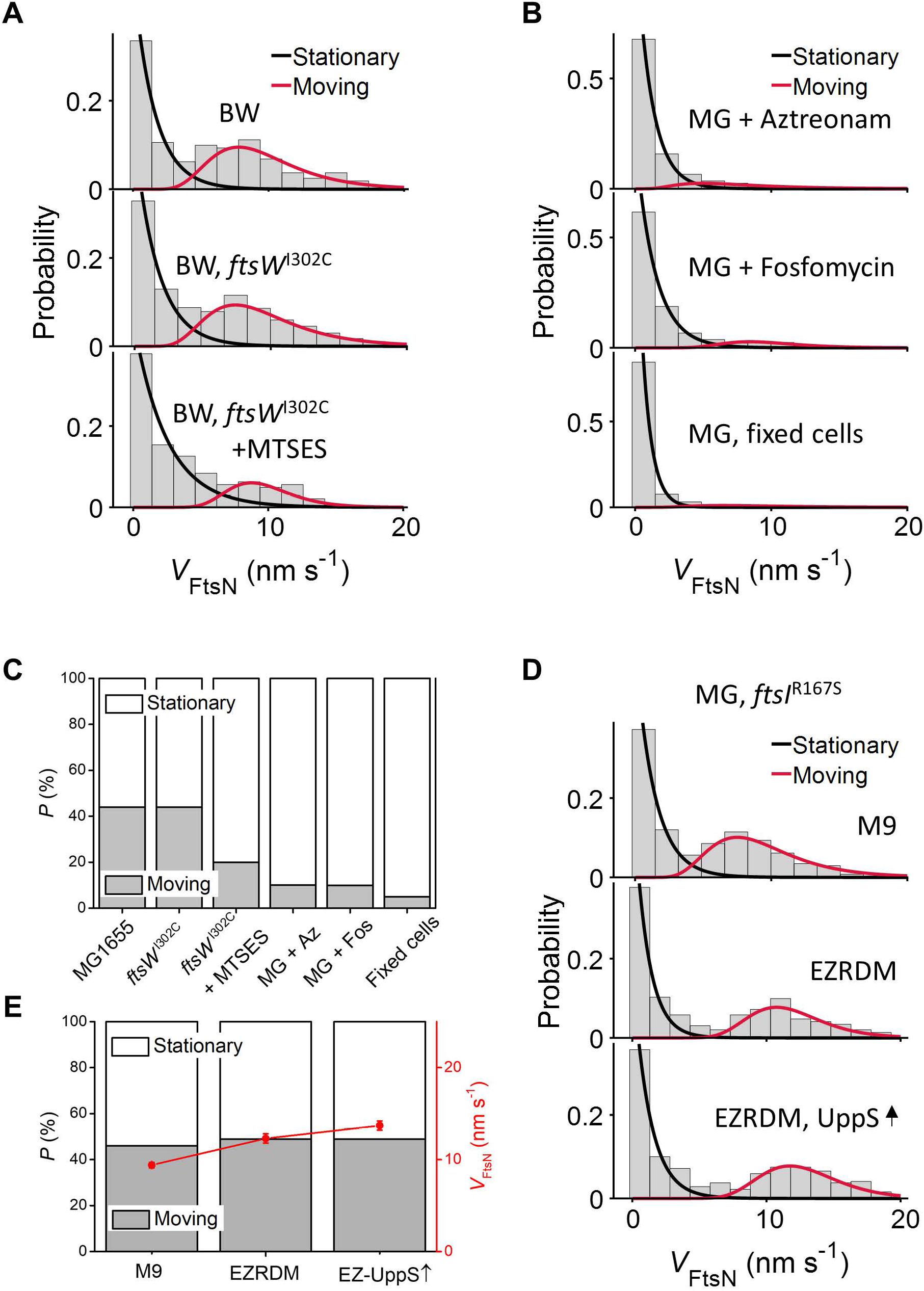
FtsN’s processive moving population is driven by sPG synthesis activity. (A) Speed distributions and the corresponding fit curves of the stationary (black) and moving (red) populations of single FtsN-Halo^SW^ molecules in the BW25113 WT (top) and *ftsW*^I302C^ variant strain in the absence (middle) or presence (bottom) of MTSES. (B) Speed distributions of single FtsN-Halo^SW^ molecules in the MG1655 WT strain treated with aztreonam (top) or fosfomycin (middle). Fixed cells without antibiotic treatment were shown as a control (bottom). (C) Percentage of the processively moving population of FtsN (grey bar) gradually decreased when sPG synthesis is inhibited under the conditions in (A and B). (D) Speed distributions of single FtsN-Halo^SW^ molecules in the MG1655 *ftsI*^R167S^ superfission variant strain background grown in M9-glucose, EZRDM or in EZRDM medium with UppS overproduction (top to bottom). (E) Percentage of the processive moving population (grey bar) and average moving speed (red cycle) under conditions in (D).

Next, we tracked the movement of FtsN-Halo^SW^ in the presence of aztreonam, an antibiotic that specifically inhibits the transpeptidase (TPase) activity of FtsI^67^. In cells treated with aztreonam (1 μg ml^-1^, 30 min), we observed that the directionally moving population of FtsN was again substantially reduced and ∼ 90% of FtsN’s SMT segments showing stationary at the septum (**Figure 3B, C, Table S10**). In addition, depleting the cell wall precursor Lipid II using fosfomycin (inhibits the essential lipid II synthesis enzyme MurA^68^, 200 μg ml^-1^, 30 min) resulted in near complete abolishment of the directionally moving population of FtsN, approaching to the background level in fixed cells (**Figure 3B, C, Table S10**). All these behaviors are, again, identical to the depletion of the slow-moving population of FtsW under identical conditions as we previously observed^42^.

To probe the dynamics of FtsN under conditions of enhanced cell wall synthesis, we made use of an *ftsI*^R167S^ superfission variant strain, which partially alleviates the need for FtsN^42^. Previously we showed that by growing *ftsI*^R167S^ cells in a rich defined medium (EZRDM) or by overexpressing the undecaprenyl pyrophosphate synthetase (UppS, an enzyme responsible for making Lipid II^69^) in the same strain, the percentage of directionally moving FtsW molecules on the slow sPG-track increased to nearly 100% and their speed increased to ∼ 13 nm s^-1^ ^42^. If FtsN is in complex with FtsWI and its movement is coupled to FtsWI’s activity, we should observe similar changes in FtsN’s dynamics.

We first tracked the dynamics of FtsN-Halo^sw^ in *ftsI*^R167S^ cells growing in minimal M9 medium and the rich defined EZRDM medium. We observed that the average speed of FtsN accelerated from 9.4 ± 0.3 nm s^-1^ in M9 to 12.3 ± 0.5 nm s^-1^ in EZRDM (**Figure 3D, E, Table S10**). Overexpressing UppS further increased the average speed of FtsN to 13.7 ± 0.5 nm s^-1^ (**Figure 3D, E, Table S10**). These increased speeds are similar to those of the slow-moving population of FtsW under the same conditions^42^. Most importantly, the distributions of the speed, processive run length and run time of FtsN-Halo^sw^ are indistinguishable from those of FtsW under the EZRDM and UppS overexpression conditions, where FtsW essentially only exhibits one slow-moving population (**Figure S11, Table S11**), strongly suggesting that FtsN forms an active, processive sPG synthesis complex with FtsWI.

### FtsN’s E domain is sufficient for forming a processive complex with FtsWI on the sPG-track

What interaction mediates the processive complex between FtsN and FtsWI? Past studies have shown that a short fragment of FtsN comprising only the second helix in the periplasmic E domain is both necessary and sufficient for cell division when overexpressed^11, 19^. An FtsN mutant containing changes in two conserved amino acids in the E domain (WYAA, with W83 and Y85 changed to alanines) fails to support cell division^19, 70^. We reason that if the E domain activates sPG synthesis by forming a processive, directional moving complex with FtsWI, the failure of the WYAA mutant to activate sPG synthesis may be mediated through the dissolution of the processive complex. Because the WYAA mutant is lethal due to the lack of FtsWI activity, to test this hypothesis, we took advantage of an *ftsB* superfission variant strain (*ftsB*^*E56A*^ *ΔftsN*) in which FtsWI is constitutively active without FtsN^19^.

We first constructed a FtsN^WYAA^-Halo^SW^ fusion and expressed it from a plasmid in the superfission variant *ftsB*^E56A^ Δ*ftsN* background (Strain JL398 in **Table S1**). As a control, we also expressed the wild-type FtsN-Halo^SW^ in the same strain background (Strain JL397 in **Table S1**). We observed that both FtsN^WYAA^-Halo^SW^ and wild-type FtsN-Halo^SW^ exhibited similar levels of midcell localization (**Figure 4A**), as FtsN’s major localization determinant—the SPOR domain—remains intact in both fusion proteins. However, the majority of FtsN^WYAA^-Halo^SW^ fusion protein remained stationary at septa as the directional moving population was significantly diminished to ∼ 11% compared to that of wild-type FtsN-Halo^SW^ (∼ 44%) (**Figure 4B, Table S12**). Combined with our previous observation that FtsW’s slow-moving population is also significantly reduced in this strain background even though FtsN is no longer essential^42^, this finding suggests that the formation of the processive sPG synthesis complex between FtsN and FtsWI is indeed mediated by the two conserved residues and crucial for activating FtsWI.

**Figure 4.**
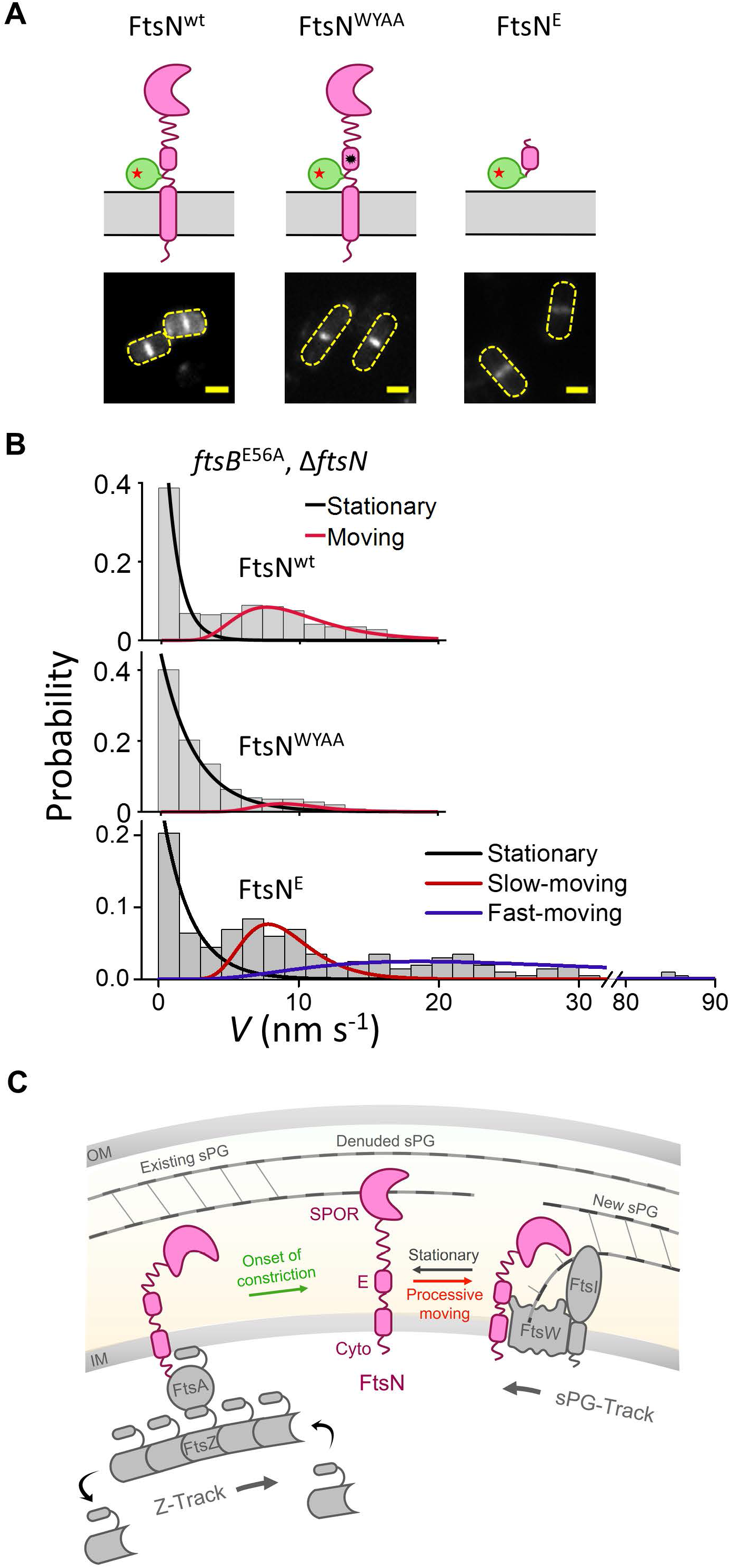
FtsN’s E domain is sufficient to form a processive complex with FtsWI on the sPG-track. (A) Schematic representation of FtsN-Halo^SW^, FtsN^WYAA^-Halo^SW^, and Halo-FtsN^E^. The black star in FtsN^WYAA^-Halo^SW^ represents the W83A and Y85A double substitution. The green circle represents the Halo tag and the red star represents the JF646 dye. Representative ensemble fluorescence cell images are shown in the bottom panel. Scale bars, 1 µm. (B) Speed distributions and corresponding fit curves for stationary (black), slow-moving (red) and fast-moving (blue) populations of single FtsN-Halo^SW^, FtsN^WYAA^-Halo^SW^, and Halo-FtsN^E^ molecules in the superfission variant *ftsB*^E56A^ Δ*ftsN* background (top to bottom). A break of the x-axis from 32 to 79 nm s^-1^ was used to accommodate the different scales of the slow- and fast-moving populations of FtsN^E^. (C) A model depicting how FtsN activates sPG synthesis. FtsN is first recruited to the septum through the interaction between its cytoplasmic tail with FtsA, and is distributed around the septum by treadmilling FtsZ polymers. Next, at the onset of constriction, FtsN binds to denuded glycan strands through its SPOR domain, which diminishes the interaction between FtsN and FtsA and renders FtsN stationary at the septum. The interaction between FtsN’s E domain with FtsWI (either directly or through FtsQLB) releases FtsN from denuded glycan strands and results the formation of an activated sPG synthesis complex, which engages in processive sPG synthesis. The active complex is sustained on the sPG-track by the presence of FtsN in the complex. Stochastic or regulated dissociation of FtsN from the complex may result in the termination of sPG synthesis, which could release FtsWI to the fast Z-track. Dissociated FtsN could rebind with denuded glycan strands, waiting for the next activation event.

Finally, to address directly whether the E domain itself is sufficient for the processive movement of FtsN, we tracked the dynamics of a Halo fusion to only the E domain containing helix 1 and the essential helix 2 (amino acids 61 to 105) in the same *ftsB*^E56A^ *∆ftsN* strain background (Strain JL399 in **Table S1**). Remarkably, although Halo-FtsN^E^ lacks the major septum localization determinant, the SPOR domain, its midcell localization is still evident (**Figure 4A**), demonstrating that its interaction with the sPG synthesis complex is independent of the SPOR domain and sufficient for its septum localization. Most interestingly, Halo-FtsN^E^ moved processively in ∼ 63% of the SMT segments (**Figure 4B**), and that a new, fast-moving population (70% of all moving segments, *v* = 28.8 ± 6.3 nm s^-1^, *μ* ± *s*.*e*.*m*., *n* = 127 segments, **Table S12**) emerged in addition to the slow-moving population (30%, *v* = 8.4 ± 1.9 nm s^-1^, *μ* ± *s*.*e*.*m*., *n* = 75 segments, **Table S12**). Because the two moving populations resemble closely the FtsZ’s treadmilling-dependent, fast-moving population and the sPG synthesis-dependent, slow-moving population of FtsW (**Figure S12**), this result strongly supports the notion that the E domain itself is sufficient to form the processive complex with FtsWI, and that such a complex can be maintained even on the Z-track when the SPOR domain is absent. In other words, the SPOR domain may be the major determinant to prevent the release of the sPG synthesis complex from the sPG-track to the Z-track.

## Discussion

FtsN, a late recruit to the *E. coli* divisome, works through FtsA and the FtsQLB complex to activate synthesis of septal PG by FtsWI. Previous work has shown that FtsWI moves directionally around the circumference of the division site on two tracks, one driven by FtsZ treadmilling (Z-track), the other driven by sPG synthesis (sPG-track). Only FtsWI in the sPG-track is actively engaged in sPG synthesis. Previous work also revealed that FtsN activates FtsWI by redistributing it from the Z-track to the sPG-track, but how FtsN does so was unclear. In principle FtsN might localize to the Z-track and prevent or even disrupt binding of FtsWI to the Z-track. Alternatively, FtsN might localize to the sPG-track and capture or retain FtsWI. Finally, FtsN might move dynamically between the two tracks with differential conformations and/or by formation of different complexes. Our findings, as detailed below, favor a model (**Figure 4C**) in which FtsN operates from the sPG-track to capture or retain FtsWI by forming a processively moving sPG synthesis complex with FtsWI and presumably other divisome proteins as well, most notably FtsQLB.

We observed that FtsN exhibits distinct septal organization and dynamics compared to those of the Z-ring. About half of the FtsN molecules in these rings are essentially static, most likely anchored by FtsN’s SPOR domain to denuded glycans in sPG. The other half move processively at a speed of ∼ 9 nm s^-1^. This velocity is essentially identical to that of the slow-moving population of FtsWI actively engaged in sPG synthesis and much slower than the ∼ 30 nm s^-1^ velocity of treadmilling FtsZ. Several additional findings support that the directionally moving population of FtsN molecules is driven by sPG synthesis on the sPG-track rather than treadmilling on the Z-track. First, the speed of FtsN is indifferent to perturbations of the treadmilling speed of FtsZ, as shown here using a series of *ftsZ* GTPase mutants (**Figure 2E**). Second, the fraction of FtsN molecules moving processively, and even their speed, can be increased by increasing the rate of sPG synthesis using a superfission mutation and increasing the supply of PG precursors (**Figure 3D**). Finally, the population of directionally moving FtsN molecules was decreased by impeding PG synthesis, which was accomplished by restricting the supply of PG precursors, inhibiting the glycosyltransferase activity of FtsW or inhibiting the transpeptidase activity of FtsI (**Figure 3A, B**).

Our findings also support that FtsN is part of an sPG synthesis complex together with FtsWI and potentially other divisome proteins such as FtsQLB. Not only the average speed of FtsN, but also its speed distribution, average run times and average run length, are, within error, identical to those of FtsWI under the EZRDM rich growth and UppS overexpression conditions (**Figure S11**). Under these two conditions, nearly all FtsW molecules are engaged in sPG synthesis and hence exhibit only one slow-moving, active population on the sPG-track, identical to that of FtsN. Moreover, tracking of various mutant derivatives of FtsN revealed that the only domain required for the processive complex formation on the sPG-track is the Essential (E) domain, which is proposed to interact with the sPG synthesis machinery FtsWI, likely via the FtsQLB complex^15, 16, 18, 19^. Most importantly, such a complex is crucial for activating and sustaining sPG synthesis in a processive manner, as a double point mutation that inactivates the E domain (WYAA) prevents formation of the processive complex and causes failure of cell division. Although the E domain has also been implicated in binding to the bifunctional PG synthases PBP1a and PBP1b^71-73^, these enzymes are not known to move processively^13, 74^, so they are not strong candidates to account for the directional movement of FtsN. They could, however, interact with the stationary population of septal FtsN, which requires further investigation.

Additionally, we have obtained evidence showing that the interaction between FtsN and FtsA, mediated by the cytoplasmic domain of FtsN, likely has a minimal contribution to the activation of FtsWI once constriction has commenced. Instead, the FtsN-FtsA interaction likely plays an important role in recruiting and redistributing FtsN along the septum through FtsZ’s treadmilling dynamics. In support of this notion, we found that abrogating the FtsN-FtsA interaction, either by deleting the cytoplasmic domain or introducing a D5N substitution, resulted in mild cell elongation, probably due to delayed recruitment of FtsN to the divisome, but did not diminish the slow-moving, sPG synthesis-engaged population of FtsN or result in cell division failure (**Figure 2F-H**). Conversely, in the absence of the periplasmic domains (E and SPOR), FtsN^Cyto-TM^ moved at the same fast speed as treadmilling FtsZ polymers and exhibited the same speed dependence in FtsZ’s GTPase mutants (**Figure 2I-K**) as we previously observed for the fast-moving population of FtsWI. These observations imply that the interaction between FtsN’s SPOR domain and denuded glycans is stronger than that between FtsN’s cytoplasmic domain and FtsA. As a consequence, full length FtsN molecules cannot end-track treadmilling FtsZ polymers once they bind to denuded glycans.

Taken together, our data suggests a model wherein FtsN activates sPG synthesis by forming a processive complex with FtsWI (**Figure 4C**). In this model, FtsN is first recruited to the septum through the interaction between its cytoplasmic tail with FtsA, and is distributed around the septum by treadmilling FtsZ polymers. This period may be too transitory for us to observe a significant population of fast-moving, full length FtsN molecules in our experiments. After the onset of constriction, FtsN binds to denuded glycan strands through its SPOR domain, which diminishes the interaction between FtsN and FtsA and creates a pool of stationary FtsN molecules at the septum. The interaction between FtsN’s E domain with FtsWI (either directly or through FtsQLB) mediates formation of an activated sPG synthesis complex that engages in processive sPG synthesis. Presumably FtsN has to release its hold on denuded glycans to move processively with FtsWI; such release might happen spontaneously or be triggered by interaction of the E domain with FtsWI. Subsequently, stochastic or regulated dissociation of FtsN from the synthesis complex may result in the termination of sPG synthesis, which could release FtsWI to the fast Z-track. Dissociated FtsN could rebind with denuded glycan strands, waiting for the next activation event. According to this model, the major function of the SPOR domain is to prevent FtsN from diffusing away from the septum or reassociating with the fast-moving FtsZ-track, which would bring FtsWI away from the sPG synthesis track. These new possibilities about FtsN’s function will be the subject of future studies.

## Acknowledgements

We thank all members of the Xiao and Weiss laboratories for helpful discussions and feedback on the manuscript, members of the Weiss lab for help with strain construction, the Microscopy Facility of Johns Hopkins School of Medicine for assistance with the TIRF-SIM imaging, and Dr. L. Lavis for sharing JF549 and JF646.

Work in the Xiao lab was supported by NIH R01GM086447 and R35GM136436 (to J.X.), GM125656 (subcontract to J.X.), a Hamilton Smith Innovative Research Award (to J.X.), and in part by NIH GM007445 (to J.W.M.). Work in the Weiss lab was supported by NIH R01GM125656 (to D.S.W.) and in part by T32AI007511 to G.M.K.

## Author contributions

D.S.W. and J.X. conceived the study. A.Y., G.M.K., D.S.W. and Z.L. constructed the strains and performed genetic and phenotypic experiments. Z.L. performed all the imaging experiments and analyzed the data. X.Y. and J.W.M. wrote the custom MATLAB script for analyzing single-molecule tracking data. Z.L. analyzed the single-molecule tracking data with help from X.Y., J.W.M. and R.M. Z.L., D.S.W. and J.X. wrote the original draft. All authors reviewed and edited the manuscript. D.S.W. and J.X. supervised the study. Funding was acquired by D.S.W. and J.X.

