## Supplementary Information for "FtsN activates septal cell wall synthesis by forming a processive complex with the septum-specific peptidoglycan synthase in *E. coli*"

### Methods

**Growth media.** Lysogeny broth (LB) was employed for routine growth and genetic manipulation of *E. coli* strains. For microscopy, cells were grown in the rich defined medium EZDRM[1] or M9-glucose minimal medium (0.4% D-glucose, 1× MEM amino acids and 1× MEM vitamin)[2]. For single-molecule tracking experiments, vitamins were omitted from the M9 medium (termed $M9^-$ minimal medium) to minimize background. Where appropriate, antibiotics were included as follows: ampicillin, 100 $\mu g\,ml^{-1}$; carbenicillin, 25 $\mu g\,ml^{-1}$; chloramphenicol, 35 $\mu g\,ml^{-1}$; kanamycin, 40 $\mu g\,ml^{-1}$; spectinomycin, 100 $\mu g\,ml^{-1}$ for plasmids and 35 $\mu g\,ml^{-1}$ for chromosomal alleles. In general antibiotics were used when propagating *E. coli* strains in LB but omitted when growing cells in minimal media for microscopy. This was possible because most of the fusions used in this study were integrated into the chromosome. However, chloramphenicol was added to minimal media for growth of strains containing the plasmid strains pJL132, pJL133, and pJL136. Where appropriate, 0.2% L-arabinose was used to express *ftsN* under $P_{BAD}$ control[3]. Isopropyl β-D-1-thiogalactopyranoside (IPTG) was used as indicated to express genes under control of modified (weakened) *Trc* promoters.

**Bacterial strains, oligonucleotide primers, and plasmids.** Standard procedures were used for analysis of DNA, PCR, electroporation, transformation, P1 transduction and integration of CRIM plasmids[4]. Bacterial strains are listed in **Table S1**, which also describes strain construction. Plasmids are described in **Table S2**, followed by descriptions of how these plasmids were made. Some plasmids were assembled by amplifying appropriate DNA fragments using Q5 DNA polymerase followed by ligation or assembly into restriction-digested vectors using T4 DNA ligase or NEBuilder HiFi DNA Assembly Master Mix, respectively (New England Biolabs). Alternatively, plasmids were assembled by amplifying appropriate insert and vector DNA fragments followed by In-Fusion cloning (Takara, In-Fusion HD Cloning Kit). The Quikchange Lightening Kit (New England Biolabs) was used for site-directed mutagenesis as needed. Oligonucleotides were from Integrated DNA Technologies (Coralville, IA), and are listed in **Table S3**. Regions of plasmids encompassing fusion genes constructed by PCR were verified by DNA sequencing.

**Purification of His6-FtsN periplasmic domain.** A fusion of a hexahistidine tag to the periplasmic domain of FtsN (residues 49-319) was overproduced in BL21(DE3) and purified on Talon affinity resin according to instructions from the manufacturer (Takara Bio USA, Inc.). Purified protein was dialyzed into 50 mM $Na_2HPO_4$, 200 mM NaCl, 5% glycerol, pH 7.5. Protein concentration was determined by BCA assay (Pierce) with BSA as standard. The yield from 1 Liter of cells was 7 mg at a concentration of 3.5 mg/ml and 80% purity as estimated from SDS-PAGE.

**Anti-FtsN anti-sera.** Rabbit anti-FtsN was raised against a maltose-binding protein fusion to the periplasmic domain of FtsN (residues 56-319) and has been described[5]. Because FtsN co-migrates in SDS-PAGE with maltose-binding protein, it was necessary to remove anti-MBP antibodies. This was accomplished by incubating the anti-serum with a concentrated cell lysate from the *E. coli* MBP overproduction strain DH5α/pMAL-c2 as described[6].

**Growth of cells for various experiments.** Cultures were grown differently depending on the experiment for which the cells were to be used. Our standard procedure to grow cells for superresolution or SMT microscopy was as follows. Starter cultures were grown overnight at 30°C or 37°C in LB, supplemented in most cases with 0.2% L-arabinose (to induce $P_{BAD}$::*ftsN*) and antibiotics if appropriate. The next day cells were washed once with M9-glucose to remove antibiotics and arabinose, then diluted 500 to 2000-fold into 3 ml M9-glucose containing IPTG as indicated to induce expression of chromosomal *ftsN* fusions; antibiotics were omitted except for plasmid strains. Cultures were incubated at room temperature with shaking until the $OD_{600}$ reached ~ 0.35 (~ 18 hours).

For the complementation assay in **Figure S2A**, overnight cultures were serially diluted into LB (no arabinose) to $OD_{600}$ = 0.1, incubated for ~ 30 min until they reached $OD_{600}$ = 0.2, then 10-fold serial dilutions were prepared in M9. Dilutions were spotted onto M9 plates, which were photographed after 18 hr incubation at 37°C. To obtain the growth curves in **Figure S2B**, overnight cultures were grown in M9-glucose. The following day, $OD_{600}$ was measured with a Nanodrop and cultures were diluted to an initial $OD_{600}$ of 0.1 in 200 µl M9-glucose in a Corning Costar sterile 96-well plate. The 96-well plates were incubated and shaken in a Tecan Infinite M200 Pro set at 30°C, with $OD_{600}$ measurements taken every 30 min for 23.5 h, shaking the plate for 3 min at 220 rpm before measuring. Doubling times were calculated from the linear phase of the log-transformed growth curve data as fitted with a straight line. For the localization experiment in **Figure S2C**, starter cultures were grown overnight at 30°C in LB supplemented with antibiotics and 0.2% L-arabinose to induce $P_{BAD}$::*ftsN*. The next day cells were washed once with M9-glucose to remove arabinose, then diluted 500 to 2000-fold into 3 ml M9-glucose containing antibiotics to select for plasmids but without IPTG (i.e., no induction was need because leaky expression from the plasmid is sufficient). Cultures were incubated at room temperature with shaking until the $OD_{600}$ reached ~ 0.35 (~ 18 hours). Cultures were grown similarly to obtain samples for the Western blots in **Figure S3**, except that antibiotics were omitted for chromosomal fusions (panels a, b, c) and IPTG was included as indicated in the figure. At the time cells were harvested for microscopy, a 0.5 ml aliquot of each culture was fixed with paraformaldehyde for cell length determinations. Cells were photographed under phase contrast and measured using Olympus cellSens Dimension software.

**In-gel assay for expression and stability of Halo fusion proteins.** Cells were grown at room temperature to $OD_{600}$ ~ 0.35 in M9-glucose with IPTG as needed. 1 ml of culture was transferred to a microfuge tube before adding 10 µl of 100 nM JF549 dye in DMSO and incubating for 30 min at room temperature in the dark. Cells were pelleted in a microfuge, the supernatant was discarded, and the pellet was taken up in ~ 70 µl 1x Laemmli Sample Buffer (LSB) to achieve a sample concentration of 5.0 $OD_{600}$ units per ml. Samples were heated for 10 min at 95°C before loading 10 µl onto a precast mini-PROTEAN TGX gel (10% polyacrylamide, from Bio-Rad, Hercules, CA). After electrophoresis the gel was washed briefly two times with distilled water, then imaged on a Sapphire Biomolecular Imager (Azure Biosystems) using the Alexa 546 setting (excitation 520 nm, emission 565 nm) with scanning parameters set to scan gel, 100-pixel resolution, slow speed.

**Western blotting.** Cells from 1 ml of culture at an $OD_{600}$ ~ 0.35 were harvested by centrifugation and the cell pellet was taken up in ~ 70 µl 1x Laemmli Sample Buffer (LSB) to achieve a sample concentration of 5.0 $OD_{600}$ units per ml. Samples were heated for 10 min at 95°C before loading 10 µl onto a precast mini-PROTEAN TGX gel (10% polyacrylamide, from Bio-Rad, Hercules, CA). Electrophoresis, transfer to nitrocellulose and blot development followed standard procedures. Primary antibody was a 1:1000 dilution of polyclonal rabbit anti-FtsN sera that had been pre-absorbed against a lysate of DH5α/pMAL-C2 as described above. Secondary antibody was horseradish peroxidase-conjugated goat anti-rabbit antibody (1:10,000; Pierce, Rockford, IL), which in turn was detected with SuperSignal WEST Pico Plus chemiluminescent substrate (Pierce, Rockford, IL). Blots were visualized with a ChemDoc Touch Imaging System (BioRad, Hercules, CA).

**Quantification of FtsN.** The wild-type strains EC251 and BW25113 were grown at room temperature in M9-glucose to $OD_{600}$ ~ 0.35 as described above. Multiple 1 ml aliquots were harvested by pelleting cells in a microfuge and taking up pellet in LSB to achieve a sample concentration of 4.5 $OD_{600}$ units per ml. Samples were pooled. In parallel, cultures were diluted and plated to determine CFUs.

To create a standard curve, purified $His_6$-FtsN periplasmic domain was mixed with an aliquot of cell extract and then serially diluted into the extract. After heating, 10 µl samples (corresponding to 1.9 x $10^7$ cells of MG1655 or 1.7 x $10^7$ cells of BW25113) were loaded onto 10% polyacrylamide gels. Subsequent steps followed the Western blotting procedures described above. Band intensities were quantified using ImageJ and used to interpolate ng of native FtsN on the blot, which was converted to number of molecules using the molecular masses of His6-FtsNperi (31,869 kDa) and native FtsN (35,793 kDa).

EC251 was determined to have on average 0.27 ng per lane. EC251 contained on average 264 molecules per cell (N = 2). BW25113 was determined to have on average 0.32 ng per lane. BW25113 contained on average 310 molecules per cell (N = 2).

**Construction of functional FtsN fusions**. FtsN has at least four functional domains spanning from the N-terminal cytoplasmic tail to the C-terminal periplasmic SPOR domain. To avoid any potential interference from the tag, we screened 11 FtsN fusions that mNeonGreen (mNeG) was fused to the N-terminus, C-terminus or inserted at internal positions of FtsN. The complementation, cell growth rates, and midcell localization images during cell division were obtained as the criteria to identify the functional fusions. Finally, the fusions with mNeG fused to the N-terminus (mNeG-FtsN) or inserted between E60 and E61 (E60-mNeG-E61) passed all the tests. The others either showed less complementation (N28-mNeG-L29), or slower growth rate (P12-mNeG-A13, N28-mNeG-L29, Q113-mNeG-L114), or polar cell localization beside the midcell localization (Q113-mNeG-L114, Q124-mNeG-M125, Q151-mNeG-T152, Q182-mNeG-T183, Q212-mNeG-T213, FtsN-mNeG), or the tag is too close to the Essential domain of FtsN (K69-mNeG-V70). For different imaging purposes, different tags or different fusions were used as indicated below.

In the 3D live-PALM imaging assay (**Figure 1B**), a mEos3.2-FtsN fusion was used since a truly monomeric photoactivatable fluorescent protein mEos3.2 [7] was needed for the single-molecule localization microscopy. Here mEos3.2 was fused to the N-terminus of FtsN.

In the FRAP assay (**Figure 1E**), a GFP-FtsN fusion was used since GFP was easily photo-bleached, which contributes a very low background signal after the photobleaching. Here GFP was fused to the N-terminus of FtsN.

In the TIRF and TIRF-SIM assays (**Figure 1F**), a mNeG-FtsN fusion was used since mNeG is much brighter and photo-stable than GFP. Here mNeG was fused to the N-terminus of FtsN.

In the SMT assay (**Figure 2-4**), a $FtsN^{E60-E61}$-Halo (termed $FtsN\text{-}Halo^{SW}$) sandwich fusion was used since E61 was included in all the FtsN mutants ($FtsN^{Cyto-TM}$ is $FtsN^{1-73}$, $FtsN^{\Delta Cyto-TM}$ is $FtsN^{61-319}$, $FtsN^{E}$ is $FtsN^{61-105}$) used in the SMT imaging. Here Halo tag was either inserted between E60 and E61, or fused to E61 on FtsN mutants. There are two advantages by using the sandwich fusion other than the N-terminal fusion in the SMT assays: (1) The Halo tag is in the same position on FtsN in all FtsN mutants, which could eliminate the potential influence caused by different positioning of the tag; (2) E60-E61 is far away from key amino acids in FtsN (e.g., D5, W83, Y85, and Q251), which could lessen potential interferences.

The functionality of mEos3.2-FtsN, GFP-FtsN, mNeG-FtsN, and $FtsN\text{-}Halo^{SW}$ fusions used in the imaging were tested by complementation, cell growth rate, cell length, and midcell localization. Their stability and expression level were tested by Western blotting (**Figure S3**).

**3D live cell SMLM imaging**. The 3D live cell SMLM imaging was conducted on a home-built microscope as previously described[8]. Briefly, the green state of mEos3.2 before activation was excited at 488-nm (laser power 40 W $cm^{-2}$) to obtain integrated green fluorescence intensity of individual cells, which was used to quantify the percentage of FtsN-ring intensity in **Figure 1D**. mEos3.2 was then photo-activated by using a 405-nm laser with intensity increased stepwise from 0 to 12 W $cm^{-2}$ to compensate for the gradually depleted pool of inactivated mEos3.2. The activated mEos3.2 was excited at 568-nm (laser power 1.6 kW $cm^{-2}$) continuously with a 10-ms exposure time.

3D imaging was achieved as previously described[8]. Briefly, a cylindrical lens (Thorlabs Inc) with 700-mm focal length was placed in the microscope emission pathway to introduce astigmatism to the single-molecule PSF[9]. TetraSpeck fluorescent microspheres with average diameter 0.1 μm (Invitrogen Molecular Probes) were used to calibrate the z-dependent changes to the shape of the astigmatic PSF. The xy positions were determined through the 2D Gaussian fitting of the PSF, while the z position was given by the calibration curve obtained by z-scanning of the fluorescent microspheres[9]. Because of the refractive index mismatch between the transmission path of the microspheres used for calibration (glass and oil) and that of the fluorescent proteins (aqueous cell environment, glass, and oil), the z values obtained from the calibration curve were rescaled by a factor of 0.75.The measurements of ring dimensions were achieved by custom MATLAB software described previously[10]. To quantitatively compare with previously reported dimensions obtained under different spatial resolutions, FtsN-ring dimensions here were deconvolved as described[10].

To quantitatively compare the distributions of clusters in FtsN- and FtsZ-rings, we used a previously established autocorrelation analysis[10]. In this analysis, all FtsN or FtsZ molecules in the ring are projected along the circumference of the ring. The spatial autocorrelation function (ACF) is calculated as the apparent probability distribution of distances between all molecule pairs. The mean ACF curve of FtsN-rings had a significantly lower correlation value at short distances (**Figure 1C**), suggesting that FtsN clusters are more homogenously distributed in FtsN-ring than those in the FtsZ-ring.

**Fluorescence recovery after photobleaching (FRAP)**. FRAP experiment was performed on a home-built microscope as previously described[11, 12]. Briefly, the excitation laser (488 nm) was split with the combination of a linear polarizer and a polarizing beam-splitting cube to generate a transmitted photobleaching beam and a reflected epifluorescence illumination beam. The transmitted beam was focused to a diffraction-limited spot on one side of the FtsN-ring for photobleaching, while the reflected beam was used to image the cell before and after photobleaching. Images were acquired every 0.5 s for 150 s after photobleaching, with a 50-ms exposure time. Custom MATLAB scripts as described previously[11] were used to analyze FRAP curves. The FtsN FRAP curve presented in **Figure 1E** and **Figure S6B** was the average of two independent experiments (~ 20 - 40 cells in each experiment). The FtsZ FRAP curve is the data from a previous work[12]. The control data in **Figure S6B** was from an adjutant cell that was not bleached in **Figure S6A** (yellow arrowhead).

The diffusion coefficient (D) of a typical inner membrane protein in prokaryotes is from 0.0075 to 0.22 $\mu m^2\ s^{-1}$, depending on the protein size, protein surface charge, and number of transmembrane helix, etc.[13]. More specifically, in our recent work, we showed that the diffusion coefficients of three divisome proteins FtsI, PBP2b, and FtsW outside the septum in wildtype *E. coli*, *B. subtilis*, and *S. pneumoniae* were 0.041, 0.038, and 0.028 $\mu m^2\ s^{-1}$, respectively[14]. The average unwrapped two-dimensional projected area of the septum in *E. coli* cells during division is ~ 0.2 $\mu m^2$ (600 nm in diameter and 100 nm in width). Half of the septal FtsN-ring was bleached in the FRAP experiment, producing a ~ 0.1 $\mu m^2$ bleaching area (A). Thus, the time that a random inner membrane divisome protein diffuses in and out of the bleaching area is ~ 2.5 s (took 0.04 $\mu m^2\ s^{-1}$ as the D and calculated by A/D). The half times of the fast recovery phase of FtsN observed in this work (2.9 ± 0.8 s) and by Söderström *et al.* (1.87 ± 0.66 s)[15] are both very close to this time, indicating that the fast recovery phase was most likely contributed by the random diffusion of FtsN molecules in and out of the bleaching area at the septum.

**TIRF and TIRF-SIM imaging and data analysis**. TIRF imaging was performed on a home-built microscope as previously described[12]. Briefly, the objective-based TIRF illumination was achieved by shifting the expanded 488-nm laser beam (Coherent Sapphire 488) off the optical axis center. The TIRF imaging angle was measured with a 20-mm right-angle prism (refractive index = 1.518, Thorlabs PS908) and fixed at ~ 70°. FtsN cluster dynamics were monitored by exciting the mNeG-FtsN fusion strain at 488 nm (laser power 0.5 W $cm^{-2}$). The exposure time was set at 1 s. 200 frames were acquired continuously without any interval dark time.

TIRF-SIM imaging was performed on a General Electric (GE) Deltavision OMX-SR super-resolution microscope with a 60X 1.49 UPlanApo oil objective and three high-speed high-sensitivity PCO sCMOS cameras to achieve higher temporal and spatial resolutions. The TIRF imaging angle was tuned from three directions by using the TetraSpeck fluorescent microspheres sample. The incident excitation power at 488-nm was adjusted to 6% transmittance (6.0% T) to minimize photobleaching. Time-lapse TIRF-SIM imaging was implemented with a 50-ms exposure time. 50 frames were acquired with 1 s interval dark time.

Cluster dynamics analysis was performed by using the ImageJ kymograph plugin (http://www.embl.de/eamnet/html/body_kymograph.html, J. Rietdorf and A. Seitz, EMBL, Heidelberg) as previously described[12]. Briefly, the fluorescence images of individual cells from TIRF or TIRF-SIM experiments were cropped, corrected for photobleaching, interpolated to 20 nm pixel$^{-1}$ via the bicubic method in ImageJ, and moving-averaged over a 4-frame window of time. The fluorescence intensity of an FtsN cluster along the direction of its movement in each frame was determined from the intensity along a line with a width of 11 interpolated pixels (~ 200 nm) manually drawn across the full length of the path of the FtsN cluster. This line was then used to plot the kymograph in **Figure 1F** and **Figure S7**. The processively moving speeds of the cluster were calculated by manually measuring the slopes of the center line of the fluorescence zigzags in the kymograph. Kymographs without obvious fluorescence zigzags were not analyzed. The speed distribution of FtsN clusters (**Figure 1G)** is the combination from TIRF and TIRF-SIM data.

The mean directional speed measured from the kymographs was at 8.8 ± 0.3 nm s$^{-1}$. The TIRF illumination region is approximal 500 nm in width according to a previous calculation[12]. Thus, the time that an FtsN cluster move across the illumination region is ~ 57 s, which is essential the same as the recovery half time of the slow phase (54 ± 10 s) we observed in the FRAP experiment, indicating that the directionally moving FtsN clusters are likely the ones contributing to the slow recovery rate of FRAP.

**Cell-labeling with Janelia Fluor 646 (JF646) dye**. Cells from 1 ml of culture at an $OD_{600}$ ~ 0.35 were harvested by centrifugation and the cell pellet was resuspended with 1 ml M9$^{-}$ minimal medium or EZRDM including 1nM JF646 (for SMT imaging) or 1µM JF646 (for ensemble imaging). The culture was mixed well with a pipette and put on a nutator at RT for 30 min. After labeling, cells were washed three times with M9$^{-}$ minimal medium or EZRDM and concentrated to 50 µl.

**SMT imaging and data analysis**. Cells were grown and labeled as described above except for certain conditions listed below. When cells were grown in EZRDM, the saturated culture was diluted 1:100 to fresh EZRDM medium with IPTG (and 0.2% L-arabinose for UppS induction) and allowed to grow at RT for 3 hr to reach the log phase. For the fixed-cell control, log-phase cells were first labeled and then fixed as described previously[11]. Cells were then loaded onto a 50 µl, 3% agarose gel pad (containing the same growth medium without antibiotic) laid in an observation chamber (FCS2, Bioptechs). The chamber was locked on the microscope stage (ASI, Eugene, OR) to minimize mechanical drifts.

For drug-treated conditions, 0.5 µl of appropriate drug solution was added to the 50 µl concentrated labeled cells. The final concentrations used were: aztreonam 1 µg $ml^{-1}$, fosfomycin 200 µg $ml^{-1}$, and MTSES 100 µM. 0.5 µl of appropriate drug solution was also added on top of the gel pad right before applying cells. The moment cells were applied was counted as time zero. With MTSES treatment, the chamber with cells was kept on the microscope stage for 60 min before the images were acquired. With aztreonam treatment, the chamber with cells was kept on the microscope stage for 30 min before the images were acquired. With fosfomycin treatment, the chamber with cells was kept on the microscope stage for 30 min before the images were acquired. All images were collected within 3 h of drug treatment.

SMT imaging was performed on an Olympus IX71 inverted microscope equipped with a 100×, 1.49 NA oil-immersion objective and Andor iXon 897 Ultra EM-CCD camera in epifluorescence-illumination mode using Metamorph 7.8.13.0 software. The focal plane was placed at ~ 250 nm from the bottom of the cell to image the molecules moving on the bottom half of the cylindrical portion of the cell body. Single molecules were tracked with 100 ms exposure time using a 647-nm laser with intensity 30 W $cm^{-2}$. 240 frames were acquired with 1Hz frame rate (1 s interval dark time between each frame), which helped to filter out molecules randomly diffusing along the cylindrical part of the cell body. 3D imaging was achieved as described above.

The data processing was similar as previously described[14, 16]. To specify, the xy positions of single molecules were determined through the 2D Gaussian fitting of the PSF with ThunderSTORM[17], a plug-in for ImageJ[18], while the z position was given by the calibration curve obtained by z-scanning of the fluorescent microspheres[9]. Because of the refractive index mismatch between the transmission path of the microspheres used for calibration (glass and oil) and that of the fluorescent proteins (aqueous cell environment, glass, and oil), the z values obtained from the calibration curve were rescaled by a factor of 0.75. A bandpass filter (60 - 300 nm) for both sigma1 and sigma2 was applied to remove the single pixel noise and out-of-focus molecules. All analysis thereafter used custom scripts in MATLAB R2020a. The localizations were linked to trajectories using a home-built MATLAB script (https://github.com/XiaoLabJHU/SMT_Unwrapping) that adopted the nearest neighbor algorithm from ref.[19]. The distance threshold was set to 300 nm per frame, which approximates to a max diffusion coefficient of ~ 0.05 $\mu m^2 s^{-1}$, or a max speed of 300 nm $s^{-1}$. To link molecules which may have blinked across frames or left the focal plane, a time threshold of 8 frames was chosen according to the off-time distribution. Only trajectories near the midpoint of the cell's long axis or near visible constriction sites where cell division takes place were used in the analysis to ensure the molecules are cell division and sPG related.

Due to the rod-shape cell envelope, the real displacement of the tracked molecules around the circumference is underestimated. The trajectories were unwrapped to one dimension using a home-built MATLAB script. We noticed that the velocity estimated from MSD curve fitting is not accurate when the dwell time of the trajectory is short (< 20 frames) or when there is more than one moving state in a single trajectory. Unwrapped trajectories were then segmented manually to determine whether a single molecule in a segment is stationary or moving processively as previously described[16]. Briefly, a segment was first fitted with a line. R, which is the ratio of the displacement and the standard deviation of fitting residuals, and P, which is the probability of processive movement, were defined and used as the criteria for the classification of segments. Through manual inspection, we determined to classify segments as processive based on a threshold of R ≤ 0.4 and P ≥ 0.5, while all others were classified as stationary. Since the confidence of classification is correlated with the segmentation length, we only consider segments longer than 5 frames to minimize classification error.

The CDF of directional moving FtsN speeds were calculated for each condition and fit to either a single or double log-normal populations:

$$CDF = P_1 \frac{\left(1 + \mathrm{erf}\left[\frac{ln\, v - u_1}{\sqrt{2\sigma_1}}\right]\right)}{2} + (1 - P_1)\frac{\left(1 + \mathrm{erf}\left[\frac{ln\, v - u_2}{\sqrt{2\sigma_2}}\right]\right)}{2}$$

Where $v$ is the moving speed for FtsN or it's mutants and $P_1$ is the percentage of the first population. For fitting with a single population, $P_1 = 1$. The values $u$ and $\sigma$ are the natural logarithmic mean and standard deviation. The average speed of each population is calculated as $\exp\left(u + \frac{\sigma^2}{2}\right)$. To estimate the error of the speed and percentage (Supplementary Tables S5), the CDF curves were bootstrapped 200 times and fit with the corresponding equation (single- or double-population).

To fit the stationary population, histograms were generated of the velocities from corresponding stationary segments in the respective FtsN condition data; the bins used in these histograms were the same used for the final respective plots as in Figures 1-4. Peak values and bin centers were used to fit a single exponential decay function:

$$f(x) = A * \exp(-\lambda x)$$

Where *A* is the amplitude of the fitted curve and $\lambda = \frac{1}{u}$. $u$ is the mean "velocity" of stationary segments.

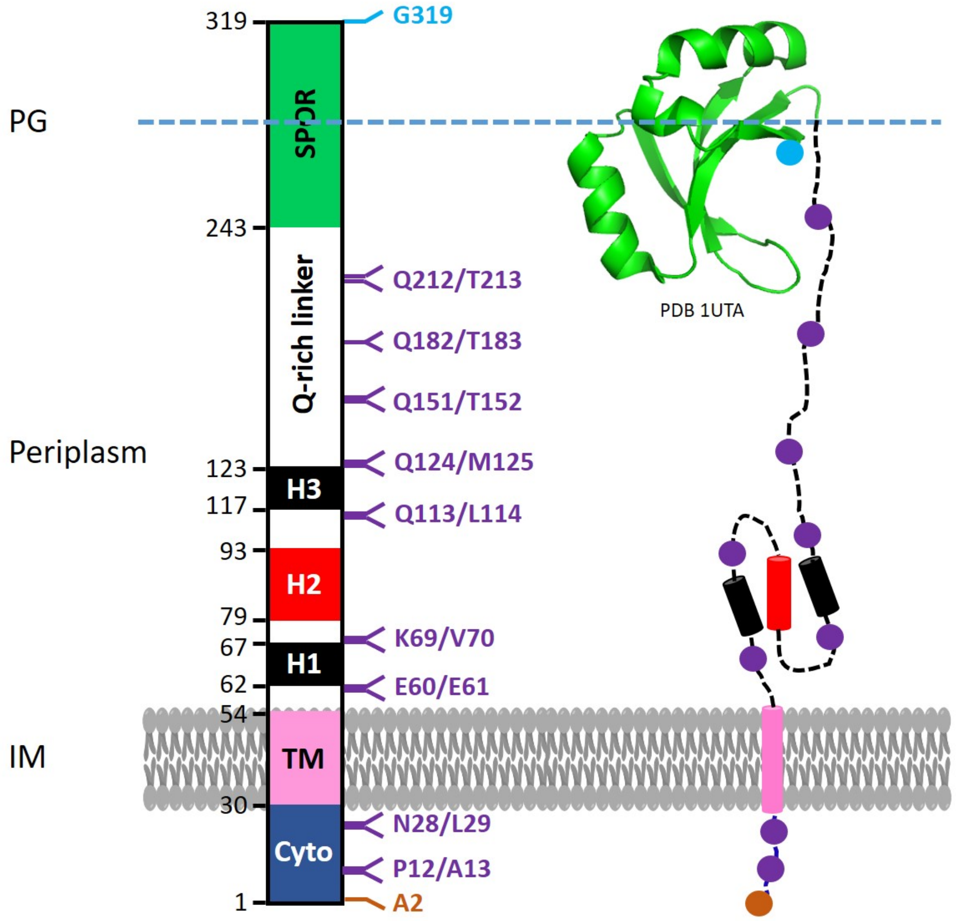


**Figure S1. Sites used for mNeG fusions to FtsN**
mNeG was fused to the N-terminus (orange), C-terminus (cyan) or inserted at internal positions (purple) of FtsN as shown by the amino acid numbers (left) and corresponding dots (right). The domain structure of FtsN is illustrated with different colors, which are the N-terminal cytoplasmic domain ($FtsN^{Cyto}$, blue), the transmembrane domain ($FtsN^{TM}$, pink), the periplasmic essential domain ($FtsN^{E}$) containing helices H1 (black), H2 (red), and H3 (black) with H2 being essential for FtsN function in cell division, and the C-terminal SPOR domain ($FtsN^{SPOR}$, green). Numbers by the left side of the domain regions refer to the amino acid range of different domains.

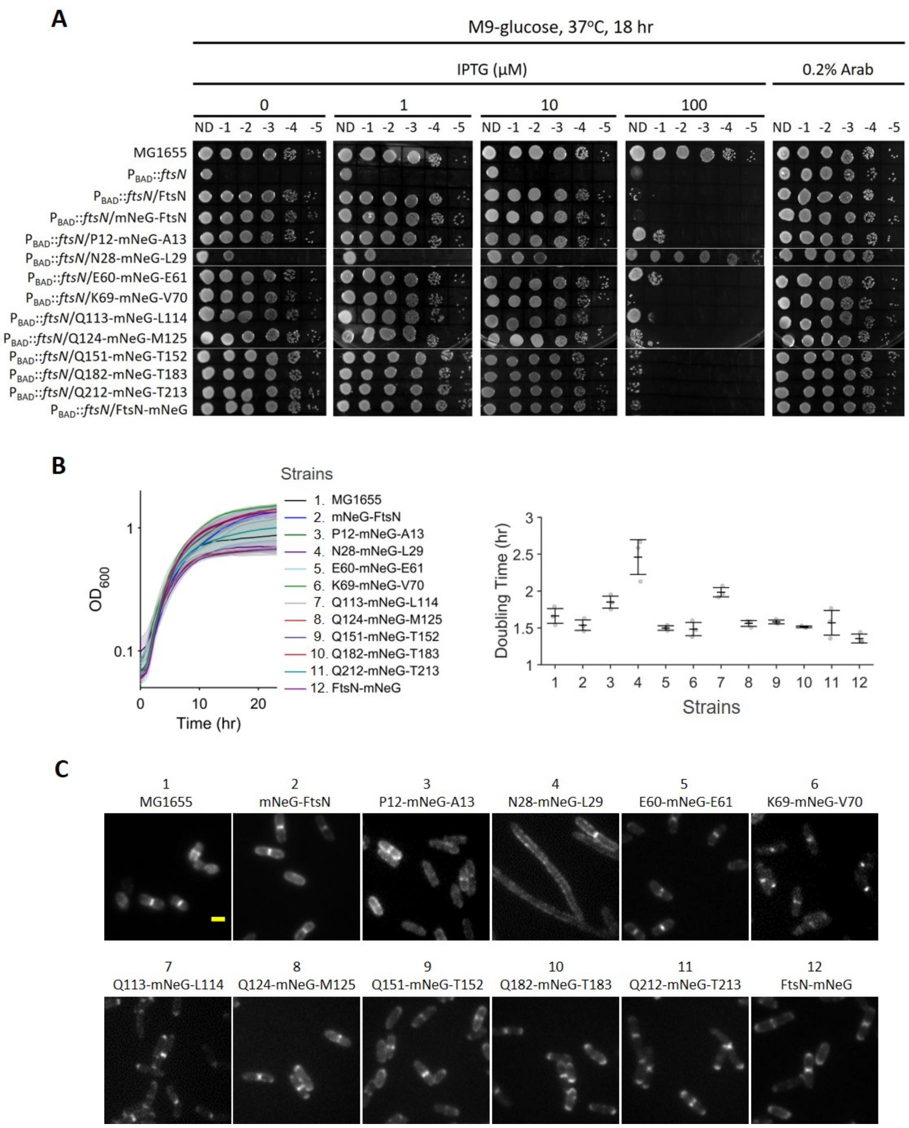


**Figure S2. Characterization of mNeG-fused FtsN constructs**

(A) To test for complementation on plates, cultures were serially diluted 10-fold, spotted onto M9 plates containing increasing IPTG concentrations, and incubated at 37°C for 18 hrs. The protein expressed under IPTG control is indicated for each strain. M9 plates containing 0.2% Arab to express chromosomal wildtype (WT) FtsN in each strain background serve as the positive control. Data were combined from two experiments. ND: no dilution.

(B) Growth curves of MG1655 and FtsN-depletion strains expressing various fusions of mNeG to FtsN in M9 minimal media at 30°C (mean ± *s.e.m.*, $n$ = 3 biological replicates). The doubling time was calculated from the growth curves (mean ± standard deviation, $n$ = 3 biological replicates). The numbers on the x-axis are the strain numbers, which correspond to the strains listed left sequentially.

(C) Septal localization of various mNeG fusions to FtsN. MG1655 cells (no FtsN fusion) were imaged using immunofluorescence staining. Cells from other strains were grown in M9 minimal media without induction. Experiment was repeated three times with similar results. Scale bar, 1 μm.

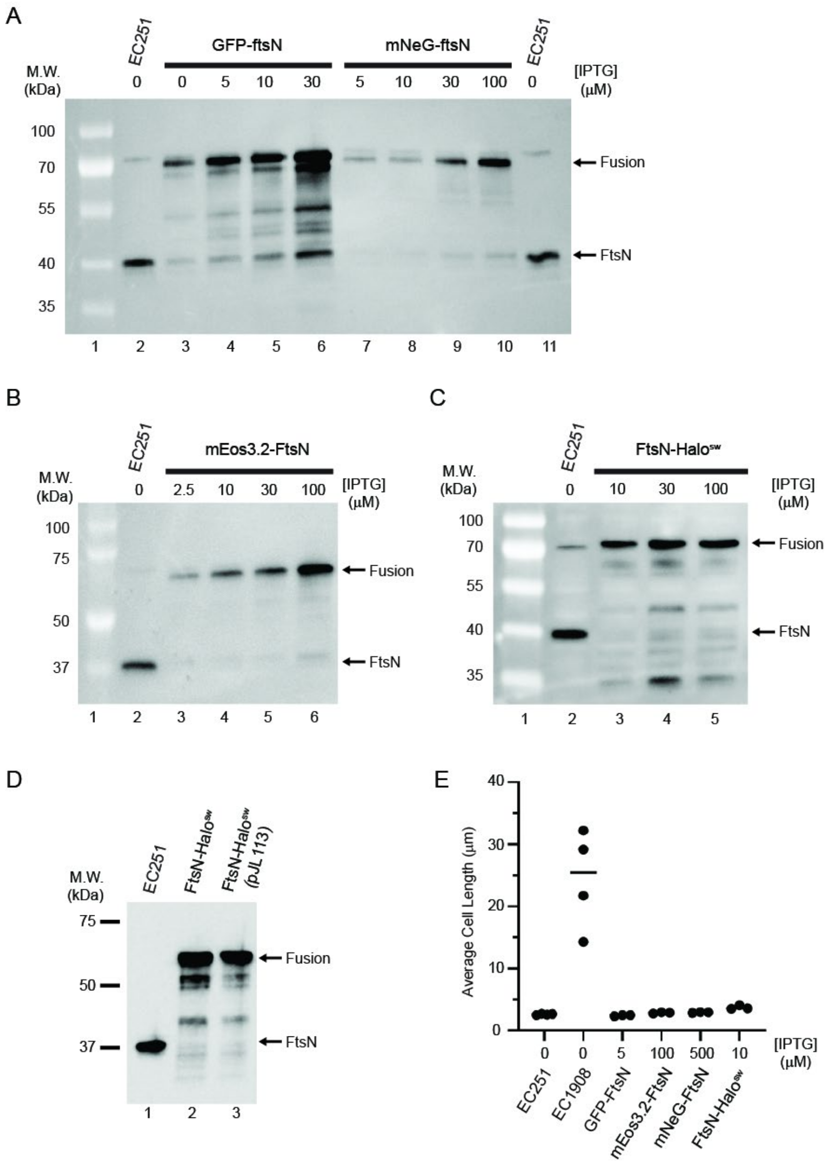


**Figure S3. Validation of FtsN fusions with mEos3.2, GFP, mNeG and Halo integrated into a chromosomal phage attachment site in an FtsN-depletion strain background (EC1908)**

(A-D) Western blots with anti-FtsN$^{peri}$ sera showing the expression levels and stability off fusion proteins at different induction conditions. For all imaging experiments, the IPTG induction level was as shown in (E). Size markers are indicated to the left of each blot. Blots are representative of at least two trials.

(E) Average cell length from 4 trials with ≥ 200 cells measured per trial. Cells were grown at room temperature in M9-glucose with IPTG as indicated to $OD_{600}$ ~0.35 before sampling for Western blotting or fixing for microscopy. The strains shown in (A-C) and (E) are EC251 (WT FtsN), EC1908 ($P_{BAD}$::*ftsN* for FtsN depletion), EC4240 (GFP-FtsN), EC4443 (mEos3.2-FtsN), EC4564 (mNeG-FtsN), and EC5234 (FtsN-Halo$^{SW}$, insertion at E60). Strains shown in (D) are EC251, EC5234, and EC5606. See more strain details in **Table S1**.

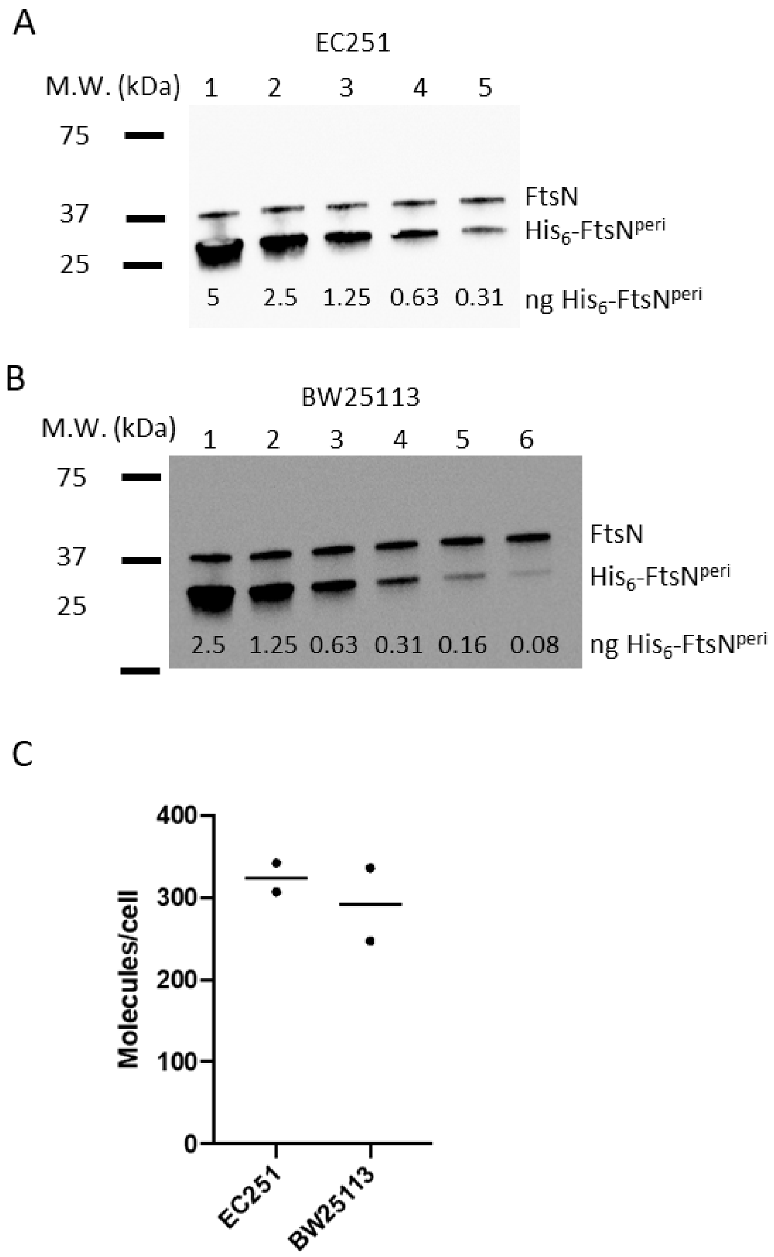


**Figure S4. Quantitation of FtsN copy number in MG1655 and BW25113 strains**
(A and B) Representative Western blots using anti-FtsN$^{peri}$ sera. The amount of FtsN in $1.9 \times 10^7$ cells of EC251 (A) and $1.7 \times 10^7$ cells of BW25113 (B) was compared to a standard curve generated by diluting purified His$_6$-FtsN periplasmic domain into the cell extracts. In the blots shown, the signal intensity of native FtsN in EC251 and BW25113 corresponded to 0.35 and 0.38 ng of His$_6$-FtsN$^{peri}$, respectively.
(C) Average number of FtsN molecules per cell (bar) in two experiments (dots) for each strain. This calculation takes into account differences in molecular mass (FtsN = 35.793 kDa, His$_6$-FtsN$^{peri}$ = 31.856 kDa) and the number of cells loaded in each lane.

**A**

$$\rho(r) = A_1\left(\frac{1}{\sqrt{\pi}\sigma_{res}}\right)e^{\frac{-r^2}{4\sigma_{res}^2}} + A_2\left(\frac{1}{\sqrt{2\pi\omega^2}}\right)e^{\frac{-(r-d_c)^2}{2\omega^2}}$$

**B**

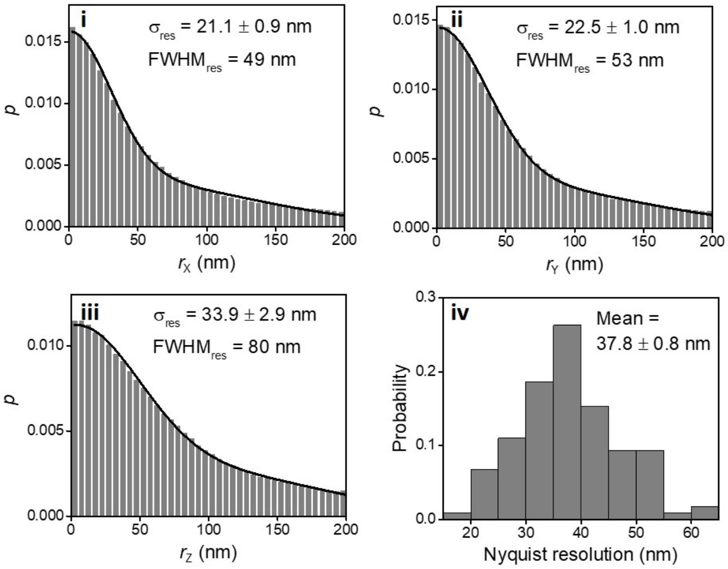


**C**

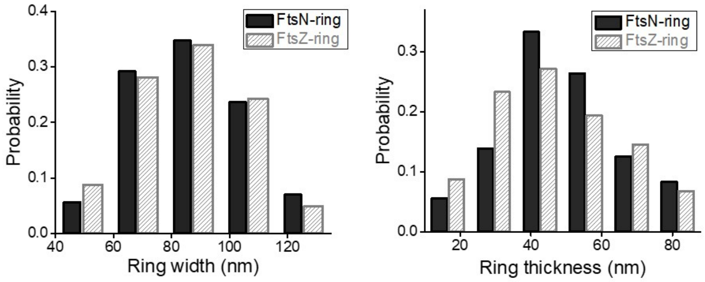


**Figure S5. Spatial resolution of 3D SMLM imaging and dimensions of FtsN-rings and FtsZ-rings**

(A) Equation describing the distribution ($p$) of pair-wise distances ($r$) between nearest neighbors in adjacent frames of live-cell SMLM data[10]. The first term represents the distribution expected for repeat observations of the same molecule with localization precision $\sigma_{res}$. The second term with Gaussian parameters $\omega$ and $d_c$ accounts for the possibility that nearest neighbors in adjacent frames may not arise from the same molecule.

(B) i-iii, Distributions of pair-wise distances between nearest neighbors in adjacent frames (gray bars) from SMLM imaging data along the x- (i), y- (ii), and z-axes (iii). Each histogram was fit using the equation in (A) to generate the black fitted curves. The achieved localization precision ($\sigma_{res}$) and spatial resolution (expressed as $FWHM_{res}$) determined from these fitted curves are displayed as insets. iv: Distribution of Nyquist resolution which was calculated as previously described[20].

(C) Distributions of resolution-deconvolved width (left) and thickness (right) of FtsN-rings (black) and FtsZ-rings (gray). There are no significant differences in the dimensions between the FtsZ- and FtsN-rings.

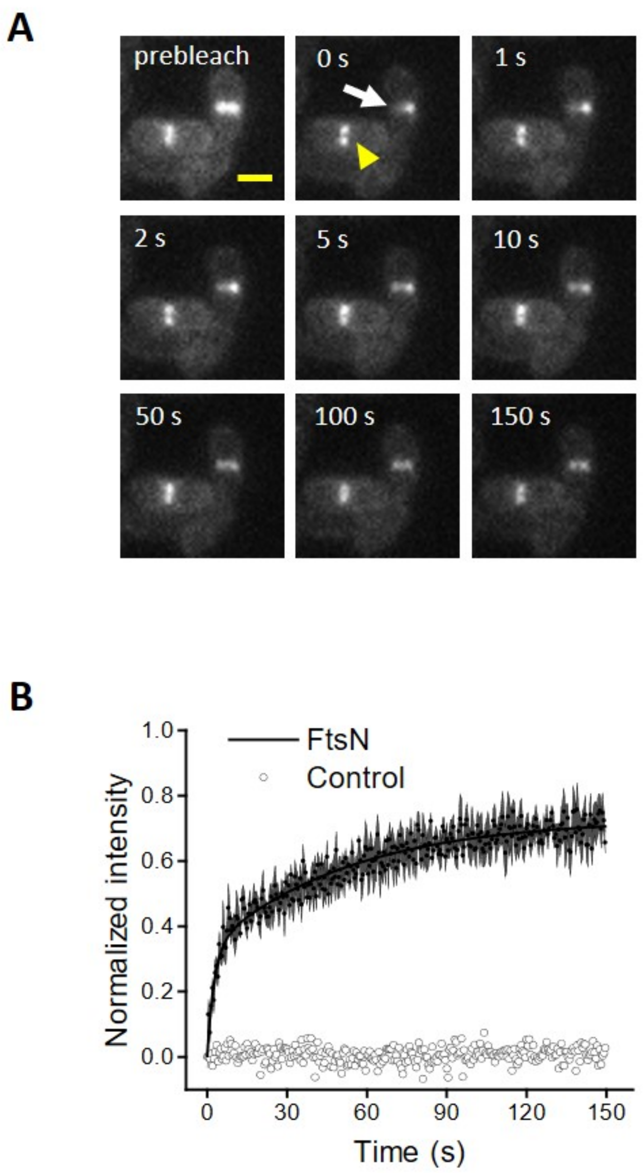


**Figure S6. FRAP analysis of FtsN**

(A) A representative FRAP imaging sequence showing the recovery of fluorescence after the photobleaching of half of the FtsN-ring (white arrow). An adjacent cell without photobleaching serves as a control (yellow arrowhead). Scale bar, 1 μm.

(B) Mean FRAP recovery curve of FtsN (black, $n$ = 58 cells, also see **Figure 1E**). The fluorescent intensity of the septum of the adjacent cell in (A) serves as a control (gray). The global photobleaching was corrected by using the fluorescent intensity outside the septum.

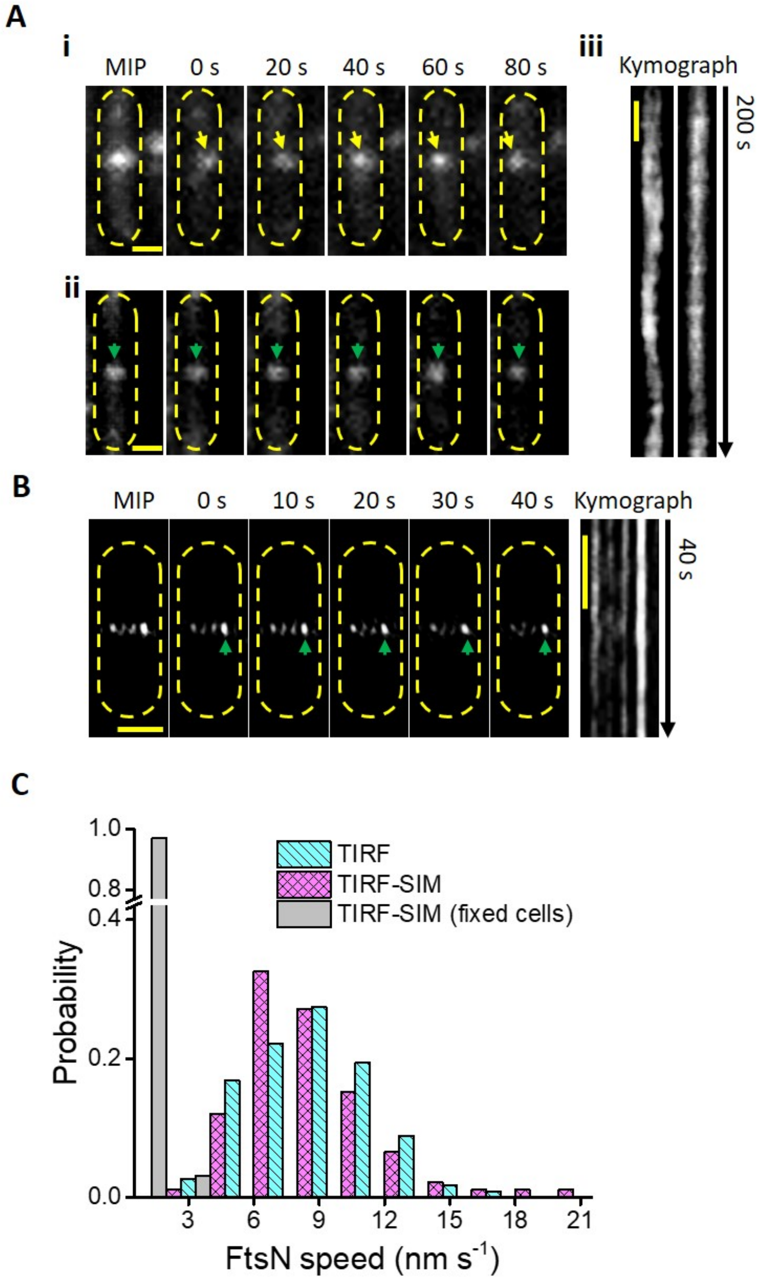


**Figure S7. FtsN clusters exhibit slow, directional motions**

(A) i-ii, Maximum intensity projection (MIP) and montages from TIRF time-lapse imaging of two cells in which a cluster is moving (i) or immobile (ii). iii, Kymographs of the cells in (i and ii) computed from the intensity along a line across the midcell are shown. Scale bars, 500 nm.

(B) Maximum intensity projection (MIP) and montages from TIRF-SIM time-lapse imaging of a fixed cell in which the clusters are immobile (left). Kymograph computed from the intensity along a line across the midcell (right). Scale bars, 500 nm.

(C) Distributions of FtsN clusters' moving speeds as measured from the kymographs.

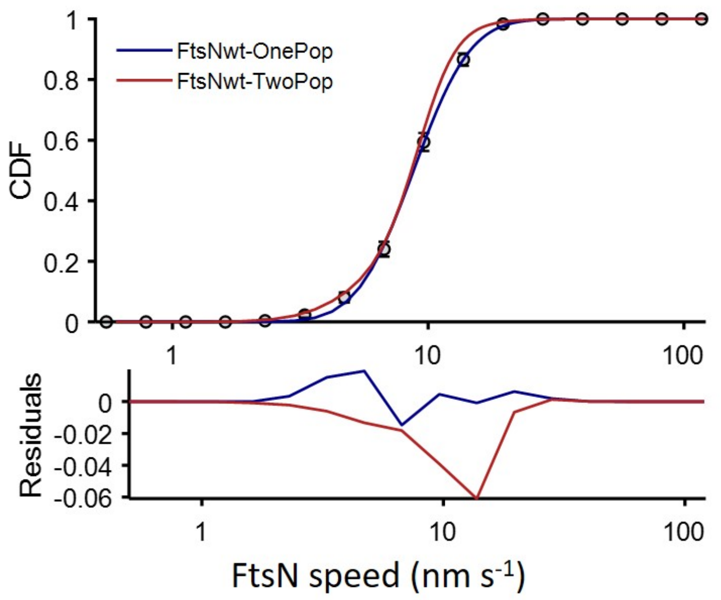


**Figure S8. Single- or double-population fitting of the cumulative probability density of FtsN's directional moving speed distribution**

CDF curve of the directional moving speed of FtsN molecules in WT MG1655 cells (black dots) was best fit by a single- (blue curve) instead of a double- (red curve) population (empirically using log-normal distribution to describe the long tail), as indicated by the residuals below. Error bars indicate *s.e.m.* from bootstrapping.

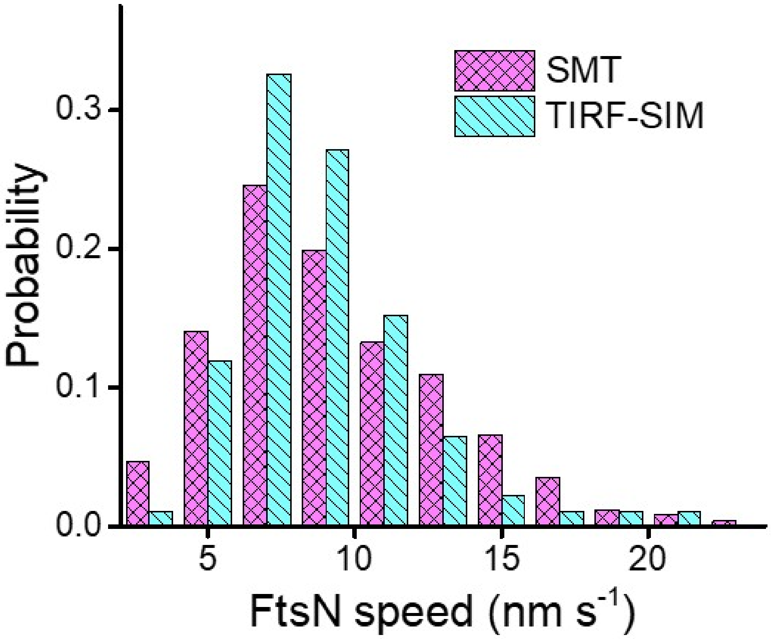


**Figure S9. Comparison of the speed distributions from SMT and TIRF-SIM**
Distributions of single FtsN molecules' moving speeds from SMT (magenta) and single FtsN clusters' moving speeds from TIRF-SIM (cyan). The *p*-value of the two-sample Kolmogorov-Smirnov (K-S) test for the two distributions is 0.15 (>0.05), indicating they are statistically identical.

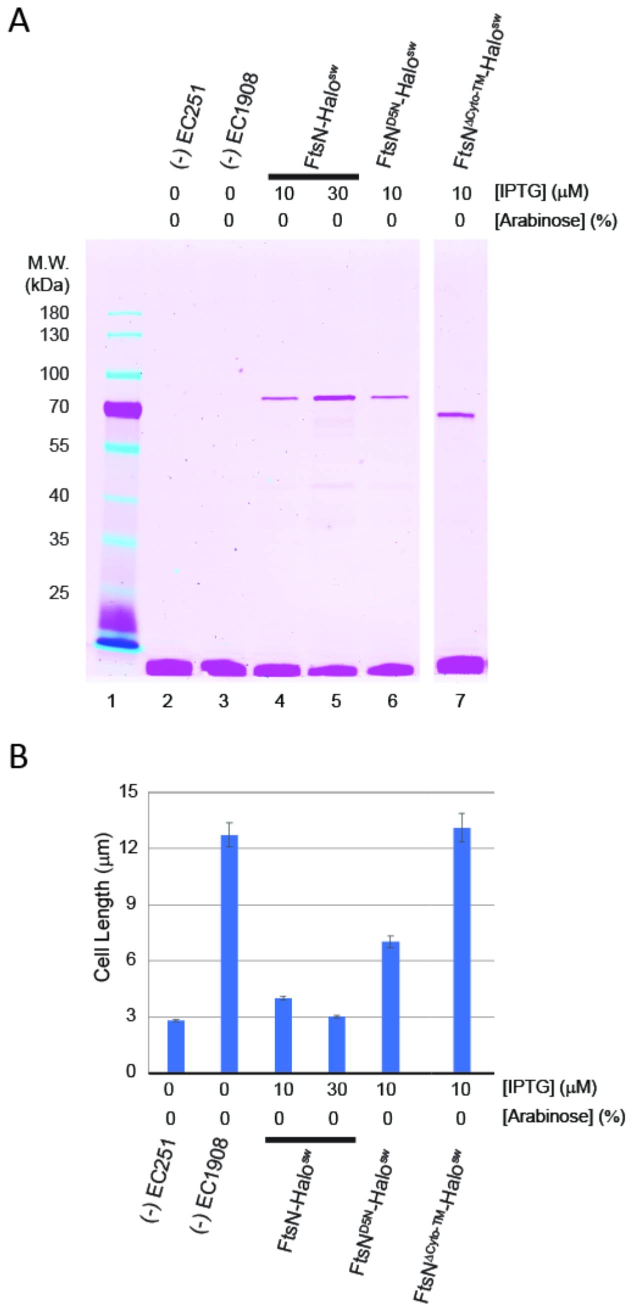


**Figure S10. Functional characterization of Halo sandwich fusions to WT and mutant FtsN proteins**

(A) Expression and stability of Halo sandwich fusions. Cells grown at room temperature in M9-glucose and IPTG as indicated were incubated with JF549 HaloTag ligand, then proteins were separated by SDS-PAGE and the gel was imaged with a fluorescence scanner. Gel is representative of two experiments.

(B) Cell length measurements. Samples from (A) were fixed and photographed under phase contrast. Data graphed as the mean and SD of two experiments. Strains shown are EC251 (WT), EC1908 ($P_{BAD}$::*ftsN*), and EC1908 derivatives that express the indicated *ftsN* fusion under control of modified *Trc* promoters (EC5234, EC5271, and EC5263, **Table S1**).

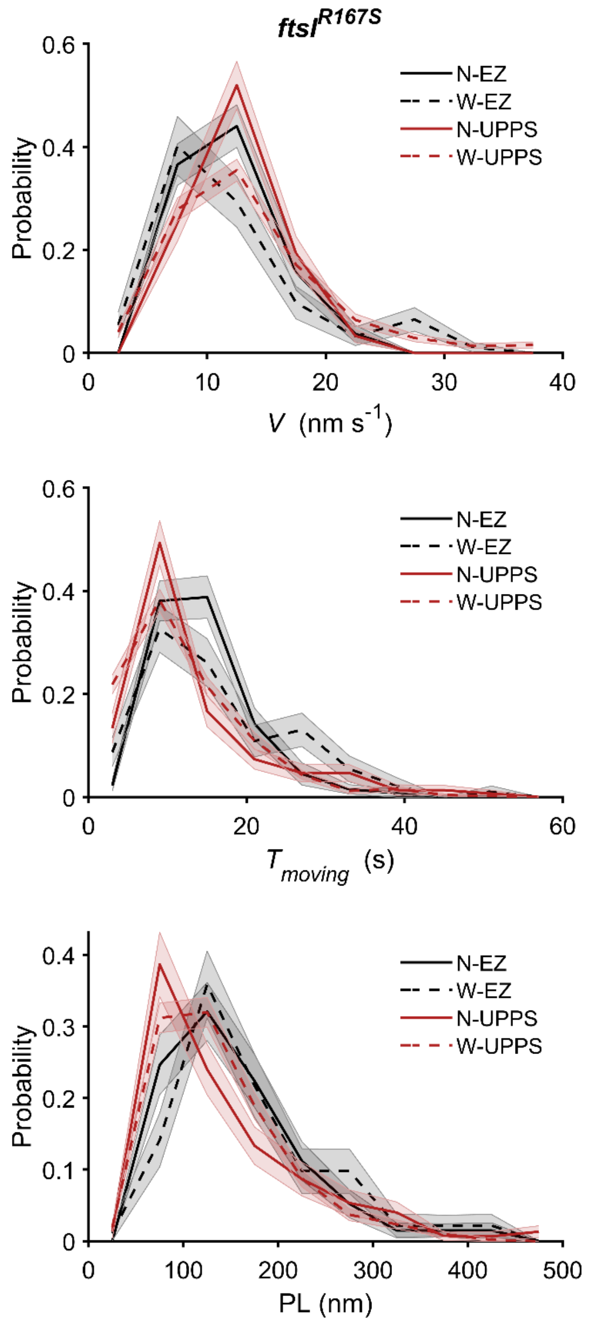


**Figure S11. Comparison of the distributions of the speed (*V*), moving dwell time (*T*_moving), and processive running length (*PL*) of FtsN and FtsW in *ftsI*$^{R167S}$ strain grown under the rich EZRDM growth condition and UppS overexpression condition**

The difference between the distributions of FtsN and FtsW under the same condition was determined to be insignificant by the two-sample Kolmogorov-Smirnov (K-S) test. The calculated *p*-values were shown in Table S11.

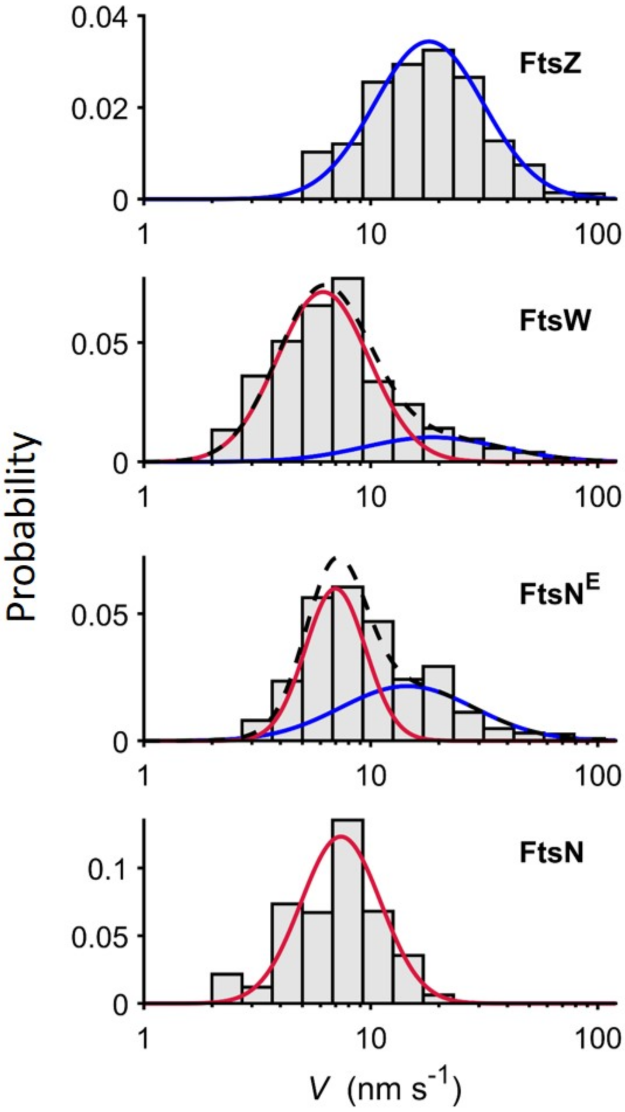


**Figure S12. Comparison of the speed distributions of FtsZ, FtsW, FtsN$^E$, and FtsN under the same growth condition**
Histograms of the speeds of FtsZ treadmilling[12], FtsW[16], FtsN$^E$, and FtsN in the log-normal scale were overlaid with one- or two-population fitting curves (slow-moving population in red, fast-moving population in blue and overall fit curve in black dashed lines).

**Table S1. Strains used in this study**

| **Strains** | **Relevant genetic markers or features** | **Source, reference or construction** |
|---|---|---|
| Strains without *ftsN* fusions | | |
| Stellar | *mcrA* Δ(*mrr-hsdRMS-mcrBC*) φ80(*lacZ*)ΔM15 Δ(*lacZYA-argF*)*U169 endA1 recA1 supE44 thi-1 gyrA96 relA1* | Takara, cloning host |
| BW25113 | *rrnB3* Δ*lacZ4787 hsdR514* Δ(*araBAD)*567 Δ(*rhaBAD)568 rph-1* | (Baba *et al.,* 2006)[21] |
| OmniMAX-2 T1R | F' [*proAB*$^{+}$ *lacI*$^{q}$ *lacZ*ΔM15 Tn*10*(Tet$^{r}$) Δ(*ccdAB*)] *mcrA* Δ(*mrr-hsdRMS-mcrBC*) φ80(*lacZ*)ΔM15 Δ(*lacZYA-argF*)*U169 endA1 recA1 supE44 thi-1 gyrA96 relA1 tonA panD* | Invitrogen, cloning host |
| XY088 | BW25113 *ftsZ*$^{E238A}$ | (Yang *et al.,* 2017)[12] |
| XY072 | BW25113 *ftsZ*$^{E250A}$ | (Yang *et al.,* 2017)[12] |
| XY089 | BW25113 *ftsZ*$^{D269A}$ | (Yang *et al.,* 2017)[12] |
| XY058 | BW25113 *ftsZ*$^{G105S}$ | (Yang *et al.,* 2017)[12] |
| EC251 | WT, Weiss lab isolate of MG1655 | (Arends & Weiss, 2004)[22] |
| EC1532 | BL21(DE3) / pDSW730 | Transformation, select Amp$^{r}$ |
| EC1908 | EC251 P$_{BAD}$::*ftsN* (Kan$^{r}$) | (Tarry *et al.,* 2009)[23] |
| JOE565 | MC4100 *ftsN*::*kan araD*+ | (Chen et al., 2001)[24] |
| BL173 | TB28 *ftsB*$^{E56A}$ *ftsN*::*kan* | (Liu *et al.,* 2015)[25] |
| PM6 | TB28 *ftsI*$^{R167S}$ | (Yang *et al.,* 2021)[16] |
| JXY559 | BW25113 *ftsW*$^{I302C}$ | (Yang *et al.,* 2021)[16] |
| JM136 | TB28 *ftsI(18-19)-Halo*$^{SW}$ | (McCausland *et al.,* 2021)[14] |
| JL273 | EC1908 / pJL098 [P$_{T5\text{-}lac}$::*ftsN* Amp$^{r}$] | This study |
| Strains with chromosomal *ftsN* fusions | | |
| EC4240 | P$_{BAD}$::*ftsN* (Kan$^{r}$) *attP*$_{HK022}$::pDSW1839 [P$_{204}$::*gfp-ftsN* Spc$^{r}$] | Integrate pDSW1839 into EC1908 using pAH69 |
| EC4443 | P$_{BAD}$::*ftsN* (Kan$^{r}$) *attP*$_{φ80}$::pDSW1890 [P$_{204_7A}$::mEos3.2-*ftsN* Spc$^{r}$] | Integrate pDSW1890 into EC1908 using pAH123 |
| EC4564 | P$_{BAD}$::*ftsN* (Kan$^{r}$) *attP*$_{φ80}$::pDSW1926 [P$_{204_7A}$::mNeG-*ftsN* Spc$^{r}$] | Integrate pDSW1926 into EC1908 using pAH123 |
| EC5230 | *attP*$_{φ80}$:: pDSW2083 [P$_{204_7A}$::*ftsN*-Halo$^{E60SW}$ Spc$^{r}$] | Integrate pDSW2083 into EC251 using pAH123 |
| EC5234 | P$_{BAD}$::*ftsN* (Kan$^{r}$) *attP*$_{φ80}$::pDSW2083 [P$_{204_7A}$::*ftsN*-Halo$^{E60SW}$ Spc$^{r}$] | Integrate pDSW2083 into EC1908 using pAH123 |
| EC5263 | P$_{BAD}$::*ftsN* (Kan$^{r}$) *attP*$_{φ80}$::pDSW2099 [P$_{204_7A}$:: *dsbA*$^{ss}$-Halo-*ftsN*$^{ΔCyto\text{-}TM}$ Spc$^{r}$] | Integrate pDSW2099 into EC1908 using pAH123 |
| EC5271 | P$_{BAD}$::*ftsN* (Kan$^{r}$) *attP*$_{φ80}$::pDSW2091 [P$_{204_7A}$::*ftsN*$^{D5N}$-Halo$^{SW}$ Spc$^{r}$] | Integrate pDSW2091 into EC1908 using pAH123 |
| EC5317 | P$_{BAD}$::*ftsN* (Kan$^{r}$) *attP*$_{φ80}$::pDSW2105 [P$_{204_7A}$::*ftsN*$^{Cyto\text{-}TM}$-Halo$^{SW}$ Spc$^{r}$] | Integrate pDSW2105 into EC1908 using pAH123 |
| EC5321 | P$_{BAD}$::*ftsN* (Kan$^{r}$) *attP*$_{φ80}$::pDSW2109 [P$_{204_7A}$::*ftsN*$^{Cyto\text{-}TM\text{-}D5N}$-Halo$^{SW}$ Spc$^{r}$] | Integrate pDSW2109 into EC1908 using pAH123 |

| | | |
|---|---|---|
| EC5333 | BW25113 *attP*$_{\varphi80}$::pDSW2083 [$P_{204\_7A}$::*ftsN*-Halo$^{E60SW}$ Spc$^{r}$] | Integrate pDSW2083 into BW25113 using pAH123 |
| EC5335 | BW25113 *ftsZ*$^{E238A}$ *attP*$_{\varphi80}$::pDSW2083 [$P_{204\_7A}$::*ftsN*-Halo$^{E60SW}$ Spc$^{r}$] | Integrate pDSW2083 into BW25113 *ftsZ*$^{E238A}$ using pAH123 |
| EC5337 | BW25113 *ftsZ*$^{E250A}$ *attP*$_{\varphi80}$::pDSW2083 [$P_{204\_7A}$::*ftsN*-Halo$^{E60SW}$ Spc$^{r}$] | Integrate pDSW2083 into BW25113 *ftsZ*$^{E250A}$ using pAH123 |
| EC5339 | BW25113 *ftsZ*$^{D269A}$ *attP*$_{\varphi80}$::pDSW2083 [$P_{204\_7A}$::*ftsN*-Halo$^{E60SW}$ Spc$^{r}$] | Integrate pDSW2083 into BW25113 *ftsZ*$^{D269A}$ using pAH123 |
| EC5341 | BW25113 *ftsZ*$^{G105S}$ *attP*$_{\varphi80}$::pDSW2083 [$P_{204\_7A}$::*ftsN*-Halo$^{E60SW}$ Spc$^{r}$] | Integrate pDSW2083 into BW25113 *ftsZ*$^{G105S}$ using pAH123 |
| EC5351 | BW25113 $P_{BAD}$::*ftsN* (Kan$^{r}$) *attP*$_{\varphi80}$::pDSW2083 [$P_{204\_7A}$::*ftsN*-Halo$^{E60SW}$ Spc$^{r}$] | P1 EC1908 x EC5333, select Kan$^{r}$ |
| EC5353 | BW25113 *ftsZ*$^{E238A}$ $P_{BAD}$::*ftsN* (Kan$^{r}$) *attP*$_{\varphi80}$::pDSW2083 [$P_{204\_7A}$::*ftsN*-Halo$^{E60SW}$ Spc$^{r}$] | P1 EC1908 x EC5335, select Kan$^{r}$ |
| EC5355 | BW25113 *ftsZ*$^{E250A}$ $P_{BAD}$::*ftsN* (Kan$^{r}$) *attP*$_{\varphi80}$::pDSW2083 [$P_{204\_7A}$::*ftsN*-Halo$^{E60SW}$ Spc$^{r}$] | P1 EC1908 x EC5337, select Kan$^{r}$ |
| EC5357 | BW25113 *ftsZ*$^{D269A}$ $P_{BAD}$::*ftsN* (Kan$^{r}$) *attP*$_{\varphi80}$::pDSW2083 [$P_{204\_7A}$::*ftsN*-Halo$^{E60SW}$ Spc$^{r}$] | P1 EC1908 x EC5339, select Kan$^{r}$ |
| EC5359 | BW25113 *ftsZ*$^{G105S}$ $P_{BAD}$::*ftsN* (Kan$^{r}$) *attP*$_{\varphi80}$::pDSW2083 [$P_{204\_7A}$::*ftsN*-Halo$^{E60SW}$ Spc$^{r}$] | P1 EC1908 x EC5341, select Kan$^{r}$ |
| EC5377 | TB28 *ftsI*$^{R167S}$ *attP*$_{\varphi80}$::pDSW2083 [$P_{204\_7A}$::*ftsN*-Halo$^{E60SW}$ Spc$^{r}$] | P1 EC5230 x PM6, select Spc$^{r}$ |
| EC5383 | BW25113 *ftsW*$^{I302C}$ *attP*$_{\varphi80}$::pDSW2083 [$P_{204\_7A}$::*ftsN*-Halo$^{E60SW}$ Spc$^{r}$] | P1 EC5230 x BW25113 *ftsW*$^{I302C}$, select Spc$^{r}$ |
| EC5387 | *ftsI*$^{R167S}$ $P_{BAD}$::*ftsN* (Kan$^{r}$) *attP*$_{\varphi80}$::pDSW2083 [$P_{204\_7A}$::*ftsN*-Halo$^{E60SW}$ Spc$^{r}$] | P1 EC1908 x EC5377, select Spc$^{r}$ |
| EC5391 | BW25113 *ftsW*$^{I302C}$ $P_{BAD}$::*ftsN* (Kan$^{r}$) *attP*$_{\varphi80}$::pDSW2083 [$P_{204\_7A}$::*ftsN*-Halo$^{E60SW}$ Spc$^{r}$] | P1 EC1908 x EC5383, select Kan$^{r}$ |
| EC5435 | BW25113 *ftsZ*$^{E238A}$ *attP*$_{\varphi8}$::pDSW2105 [$P_{204\_7A}$:: *ftsN*$^{Cyto-TM}$-Halo$^{SW}$ Spc$^{r}$] | Integrate pDSW2105 into EC5277 using pAH123 |
| EC5437 | BW25113 *ftsZ*$^{E250A}$ *attP*$_{\varphi8}$::pDSW2105 [$P_{204\_7A}$:: *ftsN*$^{Cyto-TM}$-Halo$^{E60SW}$ Spc$^{r}$] | Integrate pDSW2105 into BW25113 *ftsZ*$^{E250A}$ using pAH123 |
| EC5439 | BW25113 ftsZ$^{D269A}$ *attP*$_{\varphi8}$::pDSW2105 [$P_{204\_7A}$:: *ftsN*$^{Cyto-TM}$-Halo$^{E60SW}$ Spc$^{r}$] | Integrate pDSW2105 into BW25113 *ftsZ*$^{D269A}$ using pAH123 |
| EC5441 | BW25113 *ftsZ*$^{G105S}$ *attP*$_{\varphi8}$::pDSW2105 [$P_{204\_7A}$:: *ftsN*$^{Cyto-TM}$-Halo$^{E60SW}$ Spc$^{r}$] | Integrate pDSW2105 into BW25113 *ftsZ*$^{G105S}$ using pAH123 |

| | | |
|---|---|---|
| Strains with *ftsN* fusions on plasmids | | |
| EC5606 | EC251 $P_{BAD}$::*ftsN* (Kan$^{r}$)/pJL113 [$P_{T5-lac}$::*ftsN*-Halo$^{SW}$ Amp$^{r}$] | Transform into EC1908, select Amp$^{r}$ |
| JL397 | BL173/pJL132 [$P_{T5-lac}$::*ftsN*-Halo$^{SW}$ Cam$^{r}$] | Transform, select Cam$^{r}$ |
| JL398 | BL173/pJL133 [$P_{T5-lac}$::*ftsN*$^{WYAA}$-Halo$^{SW}$ Cam$^{r}$] | Transform, select Cam$^{r}$ |
| JL399 | BL173/pJL136 [$P_{T5-lac}$:: *dsbA*$^{ss}$-Halo-*ftsN*$^{61-105}$ Cam$^{r}$] | Transform, select Cam$^{r}$ |
| JL035 | JOE565 / pJL019 [$P_{T5-lac}$::mNeG-*ftsN* Amp$^{r}$] | Transform, select Amp$^{r}$ |
| JL247 | EC1908 / pJL107 [$P_{T5-lac}$::mNeG-*ftsN*(P12-A13)-mNeG$^{SW}$ Amp$^{r}$] | Transform, select Amp$^{r}$ |
| JL231 | EC1908 / pJL103 [$P_{T5-lac}$::mNeG-*ftsN*(N28-L29)-mNeG$^{SW}$ Amp$^{r}$] | Transform, select Amp$^{r}$ |
| JL248 | EC1908 / pJL108 [$P_{T5-lac}$::mNeG-*ftsN*(E60-E61)-mNeG$^{SW}$ Amp$^{r}$] | Transform, select Amp$^{r}$ |
| JL249 | EC1908 / pJL109 [$P_{T5-lac}$::mNeG-*ftsN*(K69-V70)-mNeG$^{SW}$ Amp$^{r}$] | Transform, select Amp$^{r}$ |
| JL250 | EC1908 / pJL110 [$P_{T5-lac}$::mNeG-*ftsN*(Q113-L114)-mNeG$^{SW}$ Amp$^{r}$] | Transform, select Amp$^{r}$ |
| JL251 | EC1908 / pJL111 [$P_{T5-lac}$::mNeG-*ftsN*(Q124-M125)-mNeG$^{SW}$ Amp$^{r}$] | Transform, select Amp$^{r}$ |
| JL232 | EC1908 / pJL100 [$P_{T5-lac}$::mNeG-*ftsN*(Q151-T152)-mNeG$^{SW}$ Amp$^{r}$] | Transform, select Amp$^{r}$ |
| JL233 | EC1908 / pJL101 [$P_{T5-lac}$::mNeG-*ftsN*(Q182-T183)-mNeG$^{SW}$ Amp$^{r}$] | Transform, select Amp$^{r}$ |
| JL234 | EC1908 / pJL102 [$P_{T5-lac}$::mNeG-*ftsN*(Q212-T213)-mNeG$^{SW}$ Amp$^{r}$] | Transform, select Amp$^{r}$ |
| JL080 | JOE565 / pJL028 [$P_{T5-lac}$::*ftsN*-mNeG Amp$^{r}$] | Transform, select Amp$^{r}$ |

### Table S2: Plasmids used in this study

| Plasmid | Relevant features/description | Source or reference |
|---|---|---|
| pAH69 | λc*I857*, *rep101*$^{ts}$ ori, $P_r$-$int_{HK022}$, $Amp^r$ | (Haldimann and Wanner, 2001)[4] |
| pAH123 | λc*I857*, *rep101*$^{ts}$ ori, $P_r$-$int_{φ80}$, $Amp^r$ | (Haldimann and Wanner, 2001)[4] |
| pAH144 | $oriR_γ$ $attP_{HK022}$ $Spc^r$, CRIM vector | (Haldimann and Wanner, 2001)[4] |
| pDSW207 | $P_{204}$::*gfp*-MCS $lacI^Q$ $Amp^r$ pBR ori | (Weiss *et al.,* 1999)[26] |
| pDSW254 | $P_{204}$::*gfp-ftsI* $lacI^Q$ $Kan^r$ pBR ori | (Weiss *et al.,* 1999)[26] |
| pDSW238 | $P_{204}$::*gfp-ftsN* $Amp^r$ pBR ori | (Jones-Carson *et al.,* 2020)[27] |
| pDSW499 | Promoterless *gfp*-MCS $oriR_γ$ $attP_{HK022}$ $Spc^r$ | (Wissel and Weiss, 2004)[5] |
| pDSW534 | $P_{204}$::*gfp*-MCS $lacI^Q$ $oriR_γ$ $attP_{HK022}$ $Spc^r$ | This study |
| pDSW984 | $P_{204}$::*gfp-dedD* $lacI^Q$ $oriR_γ$ $attP_{Φ80}$ $Spc^r$ | (Arends *et al.,* 2010)[28] |
| pDSW730 | $P_{T5/lac}$::$His_6$-$ftsN^{peri}$ $lacI^Q$ $Amp^r$ pBR ori | This study |
| pDSW1198 | $P_{204}$::*gfp* $lacI^Q$ $oriR_γ$ $attP_{Φ80}$ $Spc^r$ | (Williams *et al.,* 2013)[29] |
| pDSW1839 | $P_{204}$::*gfp-ftsN* $lacI^Q$ $oriR_γ$ $attP_{HK022}$ $Spc^r$ | This study |
| pDSW1866 | $P_{204\_7A}$::*gfp-dedD* $lacI^Q$ $oriR_γ$ $attP_{Φ80}$ $Spc^r$ | This study |
| pDSW1876 | $P_{204\_7A}$::*gfp-BamHI* $lacI^{Q*}$ $oriR_γ$ $attP_{Φ80}$ $Spc^r$ | This study |
| pDSW1884 | $P_{204\_7A}$::mEos3.2-MCS $lacI^Q$ $oriR_γ$ $attP_{Φ80}$ $Spc^r$ | This study |
| pDSW1890 | $P_{204\_7A}$::mEos3.2-*ftsN* $lacI^Q$ $oriR_γ$ $attP_{Φ80}$ $Spc^r$ | This study |
| pDSW1926 | $P_{204\_7A}$::mNeG-*ftsN* $lacI^Q$ $oriR_γ$ $attP_{Φ80}$ $Spc^r$ | This study |
| pDSW2031 | $P_{204\_7A}$::*ftsN*-$mNeG^{E60SW}$ $lacI^Q$ $oriR_γ$ $attP_{Φ80}$ $Spc^r$ | This study |
| pDSW2083 | $P_{204\_7A}$::*ftsN*-$Halo^{E60SW}$ $lacI^{Q*}$ $oriR_γ$ $attP_{Φ80}$ $Spc^r$ | This study |
| pDSW2091 | $P_{204\_7A}$::$ftsN^{D5N}$-$Halo^{E60SW}$ $lacI^{Q*}$ $oriR_γ$ $attP_{Φ80}$ $Spc^r$ | This study |
| pDSW2099 | $P_{204\_7A}$::$ftsN^{Δcyto-TM}$-$Halo^{E60SW}$ $lacI^{Q*}$ $oriR_γ$ $attP_{Φ80}$ $Spc^r$ | This study |
| pDSW2105 | $P_{204\_7A}$::$ftsN^{Cyto-TM}$-$Halo^{E60SW}$ $lacI^{Q*}$ $oriR_γ$ $attP_{Φ80}$ $Spc^r$ | This study |
| pDSW2109 | $P_{204\_7A}$::$ftsN^{Cyto-TM-D5N}$-$Halo^{E60SW}$ $lacI^{Q*}$ $oriR_γ$ $attP_{Φ80}$ $Spc^r$ | This study |
| pJC2 | $P_{BAD}$::*ftsN* $Amp^r$ pBR ori | (Chen and Beckwith, 2001)[24] |
| pXY677 | ColE1, $P_{T5-lac}$::*mNeonGreen-zapA*, $Cam^r$ | (Yang *et al.,* 2021)[16] |
| pJL015 | ColE1, $P_{T5-lac}$::*mEos3.2-ftsN*, $Amp^r$ | (Lyu *et al.,* 2016)[8] |
| pJL098 | ColE1, $P_{T5-lac}$::*ftsN*, $Amp^r$ | This study |
| pJL019 | ColE1, $P_{T5-lac}$::*mNeG-ftsN*, $Amp^r$ | This study |
| pJL107 | ColE1, $P_{T5-lac}$::*ftsN(P12-A13)-mNeG*$^{SW}$, $Amp^r$ | This study |
| pJL103 | ColE1, $P_{T5-lac}$::*ftsN(N28-L29)-mNeG*$^{SW}$, $Amp^r$ | This study |
| pJL108 | ColE1, $P_{T5-lac}$::*ftsN(E60-E61)-mNeG*$^{SW}$, $Amp^r$ | This study |

| | | |
|---|---|---|
| pJL109 | ColE1, $P_{T5\text{-}lac}$::*ftsN(K69-V70)-mNeG^SW^*, $Amp^r$ | This study |
| pJL110 | ColE1, $P_{T5\text{-}lac}$::*ftsN(Q113-L114)-mNeG^SW^*, $Amp^r$ | This study |
| pJL111 | ColE1, $P_{T5\text{-}lac}$::*ftsN(Q124-M125)-mNeG^SW^*, $Amp^r$ | This study |
| pJL100 | ColE1, $P_{T5\text{-}lac}$::*ftsN(Q151-T152)-mNeG^SW^*, $Amp^r$ | This study |
| pJL101 | ColE1, $P_{T5\text{-}lac}$::*ftsN(Q182-T183)-mNeG^SW^*, $Amp^r$ | This study |
| pJL102 | ColE1, $P_{T5\text{-}lac}$::*ftsN(Q212-T213)-mNeG^SW^*, $Amp^r$ | This study |
| pJL028 | ColE1, $P_{T5\text{-}lac}$::*ftsN-mNeG*, $Amp^r$ | This study |
| pJL113 | ColE1, $P_{T5\text{-}lac}$::*ftsN(E60-E61)-Halo^SW^*, $Amp^r$ | This study |
| pXY027 | ColE1, $P_{T5\text{-}lac}$::*ftsZ-GFP*, $Cam^r$ | (Buss *et al.,* 2015)[11] |
| pJL119 | ColE1, $P_{T5\text{-}lac}$::*ftsN*, $Cam^r$ | This study |
| pJL132 | ColE1, $P_{T5\text{-}lac}$::*ftsN(E60-E61)-Halo^SW^*, $Cam^r$ | This study |
| pJL133 | ColE1, $P_{T5\text{-}lac}$::*ftsN(E60-E61, WYAA)-Halo^SW^*, $Cam^r$ | This study |
| pJL136 | ColE1, $P_{T5\text{-}lac}$::*$dsbA^{ss}$-Halo-$ftsN^{61\text{-}105}$*, $Cam^r$ | This study |
| pCH650 | pACYC*, cat araC* $P_{BAD}$::*uppS* | (Yang *et al.,* 2021)[16] |

pDSW534. A 1706 bp fragment encoding *$lacI^Q$* and $P_{204}$::*gfp* was obtained by digesting pDSW254 with NdeI, SphI and XmnI. The fragment was ligated into pDSW499 after digestion with NdeI and SphI.

pDSW730. Amplify periplasmic domain of *ftsN* with primers P760 and P761. The 833 bp fragment was digested with BamHi and HindIII, then ligated into the same sites of pQE-80L (Qiagen).

pDSW1839. Amplify *gfp-ftsN* from pDSW238 with primers P2108 and P2109. The 1142 bp product was digested with MfeI and SacI, then ligated into the same sites of pDSW534.

pDSW1866. The $P_{204}$ promoter in pDSW984 was replaced with a weaker $P_{204\_7A}$ promoter using isothermal assembly to insert a 675 bp gBlock into the SfoI and NdeI sites of pDSW984.

pDSW1876. The $P_{204}$ promoter in pDSW1198 was replaced with a weaker $P_{204\_7A}$ promoter using isothermal assembly to insert a 675 bp gBlock into the SfoI and NdeI sites of pDSW1198. The *$lacI^Q$* allele is designated *$lacI^{Q*}$* because it was later found to have a frame shift mutation, a deletion of T999. The last 28 amino acids of wild-type LacI (NTQTASPRALADSLMQLARQVSRLESGQ) become KRKPPLPARWPIH. The mutant Lac repressor is still active but does not repress as well as wild-type, so leaky expression is about two-fold higher.

pDSW1884. Amplify mEos3.2 from pJL015 with primers P2163 and P2164. The 757 bp fragment was digested with AflII and MfeI, then ligated into AflII-EcoRI digested pDSW1866.

pDSW1890. Amplify *ftsN* from pJL015 with P2178 and P2179. The 1029 bp product was digested with EcoRI and BamHI, and ligated into the same sites of pDSW1884.

pDSW1926. Amplify mNeonGreen from pJL019 with primers P2222 and P2223. The 744 bp product was digested with AflII and EcoRI, then ligated into the same sites of pDS1890.

pDSW2031. Amplify the *ftsN*(E60-mNeG) sandwich fusion from pJL108 with P2392 and P2393. The 1730 bp product was digested with AflII and BamHI, then ligated into the same sites of pDSW1876.

pDSW2083. Constructed from a four-fragment Gibson Assembly. The vector backbone was obtained by digestion of pDSW2035 with AflII and KpnI. The inserts were a 231 bp fragment amplified from pDSW2031 with primers P2439 and P2445, an 891 bp fragment amplified from pDSW2035 with primers P2446 and P2447, and an 804 bp fragment amplified from pDSW2031 with primers P2448 and P2449.

pDSW2091. Amplify the *ftsN* (E60-Halo) sandwich fusion from pDSW2083 using primers P2422 and P2460. P2422 introduces a D5N amino acid substitution. The 2009 bp product was digested with AflII and KpnI, then ligated into the same sites of pDSW2035.

pDSW2099. Obtain a 946 bp $dsbA^{ss}$-Halo fragment from pJL074 by digestion with AflII and AccI. This fragment was ligated into the same sites of pDSW2083.

pDSW2105. Amplify a fragment of the *ftsN*(E60-Halo) sandwich fusion from pDSW2083 with primers P2182 and P2525. The 1357 bp PCR product encodes *ftsN* residues 1-73, a Halo tag inserted between *ftsN* residues 60-61, followed by *ftsN* residues 61-73. This DNA fragment was digested with AflII and KpnI, then ligated into the same sites of pDSW2035.

pDSW2109. Amplify part of the $ftsN^{D5N}$ (E60-Halo) sandwich fusion from pDSW2091 using primers P2182 and P2525. The 1357 bp product was digested with AflII and KpnI, then ligated into the same sites of pDSW2035.

pJL098. Amplify the vector backbone from pJL015 ($P_{T5\text{-}lac}$::*mEos3.2-ftsN*)[8] with primers 13 and 72. Amplify *ftsN* gene from pJL015 with primers 127 and 128. The two DNA fragments were then joined by the In-Fusion Cloning Kit to generate plasmid pJL098 ($P_{T5\text{-}lac}$::*ftsN*, $Amp^r$).

pJL019. Amplify *ftsN* gene with the vector backbone from pJL015 with primers 13 and 72. Amplify *mNeonGreen* gene from pXY677 ($P_{T5\text{-}lac}$::*mNeonGreen-zapA*)[16] with primers 11 and 12. The two DNA fragments were then joined by the In-Fusion Cloning Kit to generate plasmid pJL019 ($P_{T5\text{-}lac}$::*mNeG-ftsN*).

pJL107. Amplify $ftsN^{12\text{-}13}$ gene with the vector backbone from pJL098 with primers 148 and149. Amplify *mNeonGreen* gene from pJL019 with primers 150 and 151. The two

DNA fragments were then joined by the In-Fusion Cloning Kit to generate plasmid pJL107 ($P_{T5\text{-}lac}$::*ftsN(P12-A13)-mNeG*$^{SW}$).

pJL103. Amplify *ftsN*$^{28\text{-}29}$ gene with the vector backbone from pJL098 with primers 141 and142. Amplify *mNeonGreen* gene from pJL019 with primers 143 and 144. The two DNA fragments were then joined by the In-Fusion Cloning Kit to generate plasmid pJL103 ($P_{T5\text{-}lac}$::*ftsN(N28-L29)-mNeG*$^{SW}$).

pJL108. Amplify *ftsN*$^{60\text{-}61}$ gene with the vector backbone from pJL098 with primers 152 and153. Amplify *mNeonGreen* gene from pJL019 with primers 154 and 155. The two DNA fragments were then joined by the In-Fusion Cloning Kit to generate plasmid pJL108 ($P_{T5\text{-}lac}$::*ftsN(E60-E61)-mNeG*$^{SW}$).

pJL109. Amplify *ftsN*$^{69\text{-}70}$ gene with the vector backbone from pJL098 with primers 156 and157. Amplify *mNeonGreen* gene from pJL019 with primers 158 and 159. The two DNA fragments were then joined by the In-Fusion Cloning Kit to generate plasmid pJL109 ($P_{T5\text{-}lac}$::*ftsN(K69-V70)-mNeG*$^{SW}$).

pJL110. Amplify *ftsN*$^{113\text{-}114}$ gene with the vector backbone from pJL098 with primers 160 and161. Amplify *mNeonGreen* gene from pJL019 with primers 162 and 163. The two DNA fragments were then joined by the In-Fusion Cloning Kit to generate plasmid pJL110 ($P_{T5\text{-}lac}$::*ftsN(Q113-L114)-mNeG*$^{SW}$).

pJL111. Amplify *ftsN*$^{151\text{-}152}$ gene with the vector backbone from pJL098 with primers 164 and165. Amplify *mNeonGreen* gene from pJL019 with primers 166 and 167. The two DNA fragments were then joined by the In-Fusion Cloning Kit to generate plasmid pJL111 ($P_{T5\text{-}lac}$::*ftsN(Q124-M125)-mNeG*$^{SW}$).

pJL100. Amplify *ftsN*$^{124\text{-}125}$ gene with the vector backbone from pJL098 with primers 129 and130. Amplify *mNeonGreen* gene from pJL019 with primers 131 and 132. The two DNA fragments were then joined by the In-Fusion Cloning Kit to generate plasmid pJL100 ($P_{T5\text{-}lac}$::*ftsN(Q151-T152)-mNeG*$^{SW}$).

pJL101. Amplify *ftsN*$^{182\text{-}183}$ gene with the vector backbone from pJL098 with primers 133 and134. Amplify *mNeonGreen* gene from pJL019 with primers 135 and 136. The two DNA fragments were then joined by the In-Fusion Cloning Kit to generate plasmid pJL101 ($P_{T5\text{-}lac}$::*ftsN(Q182-T183)-mNeG*$^{SW}$).

pJL102. Amplify *ftsN*$^{212\text{-}213}$ gene with the vector backbone from pJL098 with primers 137 and138. Amplify *mNeonGreen* gene from pJL019 with primers 139 and 140. The two DNA fragments were then joined by the In-Fusion Cloning Kit to generate plasmid pJL102 ($P_{T5\text{-}lac}$::*ftsN(Q212-T213)-mNeG*$^{SW}$).

pJL028. Amplify *ftsN* gene with the vector backbone from pJL015 with primers 13 and 72. Amplify *mNeonGreen* gene from pJL019 with primers 39 and 40. The two DNA fragments were then joined by the In-Fusion Cloning Kit to generate plasmid pJL028 ($P_{T5\text{-}lac}$::*ftsN-mNeG*).

pJL113. Amplify *ftsN*$^{60\text{-}61}$ gene with the vector backbone from pJL098 with primers 152 and153. Amplify *Halo* gene from the chromosome of strain JM136 which contains the sandwich *Halo-ftsI* gene[14] with primers 177 and 178. The two DNA fragments were then joined by the In-Fusion Cloning Kit to generate plasmid pJL113 ($P_{T5\text{-}lac}$::*ftsN(E60-E61)-Halo*$^{SW}$, Amp$^r$).

pJL119. Amplify the vector backbone from pXY027 ($P_{T5\text{-}lac}$::*ftsZ-GFP*, Cam$^r$)[11] with primers 72 and 126. Amplify *ftsN* gene from pJL015 with primers 127 and 128. The two DNA fragments were then joined by the In-Fusion Cloning Kit to generate plasmid pJL119 ($P_{T5\text{-}lac}$::*ftsN*, Cam$^r$).

pJL132. Amplify *ftsN*$^{60\text{-}61}$ gene with the vector backbone from pJL119 with primers 152 and153. Amplify *Halo* gene from the chromosome of strain JM136 which contains the sandwich *Halo-ftsI* gene with primers 177 and 178. The two DNA fragments were then joined by the In-Fusion Cloning Kit to generate plasmid pJL132 ($P_{T5\text{-}lac}$::*ftsN(E60-E61)-Halo*$^{SW}$, Cam$^r$).

pJL133. The pJL133 ($P_{T5\text{-}lac}$::*ftsN(E60-E61, WYAA)-Halo*$^{SW}$, Cam$^r$) plasmid was constructed from the pJL132 plasmid using the QuikChange protocol (Agilent) with the primers 60 and 61 to mutate the nucleotide sequence encoding for W83A, Y85A.

### Table S3. Oligonucleotides used in this study

| Name | Sequence (5'->3') | Comment |
|---|---|---|
| P760 | CGGGATCCGGTGGTCTGTACTTCATTACG | Forward primer to clone *ftsN* periplasmic domain into pQE-80L |
| P761 | CCCAAGCTTTCAACCCCCGGCGGCGAGCCG | Reverse primer to clone *ftsN* periplasmic domain into pQE-80L |
| P2108 | CTCCAATTGGCGATGGCCCTGTCCT | Forward primer to clone *gfp-ftsN* into pDSW534 |
| P2109 | GATGAGCTCTCAACCCCCGGCGGCGAG | Reverse primer to clone *gfp-ftsN* into pDSW534 |
| P2163 | CAGCTTAAGACACAGGAAACAGACCATGAGTGCGATTAAGCCAG | Forward primer to clone mEos3.2 into pDSW1866 |
| P2164 | CTGCAATTGCTGCAGGTCGACTCTAGAGGATCCCCGGGTACCGAGCTCGAATTCTCGTCTGGCATTGTCAGG | Reverse primer to clone mEos3.2 into pDSW1866 |
| P2178 | CCAGAATTCATCAACAAGTTTGTACAAAAAAGCAGGCTC | Forward primer to clone *ftsN* into pDSW1884 |
| P2179 | TGGGGATCCTCAACCCCCGGCGGCGAG | Reverse primer to clone *ftsN* into pDSW1884 |
| P2182 | CTGTCTACTCTGGAGATTTCCGGTGGTGGCGGTGGTAGTGCGGAGAAAAAGACGAACGC | Forward primer for cloning *ftsN*$^{\text{Cyto-TM}}$ *and ftsN*$^{\text{D5N-Cyto-TM}}$ into pDSW2035 |
| P2222 | CAATTCTTAAGACACAGGAAACAGACCATGGTGAGCAAAGGCGAAGAAG | Forward primer to clone mNeonGreen into pDSW1890 |
| P2223 | GTGGAATTCTTTATACAGTTCATCCATGCCCATC | Reverse primer to clone mNeonGreen into pDSW1890 |
| P2392 | CACCTTAAGACACAGGAAACAGACC ATGGCACAACGAGATTATGTAC | Forward primer to clone *ftsN* sandwich fusions into pDSW1876 |
| P2393 | CACGGGATCCTTAACCCCCGGCGG | Reverse primer to clone *ftsN* sandwich fusions into pDSW1876 |
| P2394 | CACGAGATCTTTAACCCCCGGCGG | Alternative reverse primer for cloning sandwich fusions into pDSW1876 |
| P2422 | CAATTCTTAAGACACAGGAAACAGACCATGGCACAACGAAATTATGTACGCCG | Forward primer for cloning *ftsN*$^{\text{D5N}}$-Halo$^{\text{SW}}$ into pDSW2035; introduces D5N substitution |
| P2439 | GAGCGGATAACAATTCTTAAGACACAGGAAACAGACCATGGC | Forward primer for constructing *ftsN-Halo*$^{SW}$ plasmid |

| | | |
|---|---|---|
| P2445 | TTTCGGATCCGCTACCACCGCCACCTTC | Reverse primer for constructing *ftsN-Halo*$^{SW}$ plasmid |
| P2446 | CGGTGGTAGCGGATCCGAAATCGGTACTGGCT | Forward primer for constructing *ftsN-Halo*$^{SW}$ plasmid |
| P2447 | CACCGCCACCACCGGAAATCTCCAGAGTAGACAGC | Reverse primer for constructing *ftsN-Halo*$^{SW}$ plasmid |
| P2448 | GATTTCCGGTGGTGGCGGTGGTAGCGAGTCC | Forward primer for constructing *ftsN-Halo*$^{SW}$ plasmid |
| P2449 | AACATGAGAATTCGAGCTCGGTACCCGGGGATCCTTAACCCCCG | Reverse primer for constructing *ftsN-Halo*$^{SW}$ plasmid |
| P2460 | CTAGAGGATCCCGTGGAAAAATGTGACTTTTATCAC | Forward primer for cloning *ftsN*$^{D5N}$-Halo$^{SW}$ into pDSW2035 |
| P2525 | CTCGGTACCCGGGGATCCTTAGTTTCCGGTCACTTTCTGGCTTTG | Reverse primer for cloning *ftsN*$^{Cyto-TM}$ *and ftsN*$^{D5N-Cyto-TM}$ into pDSW2035 |
| 13 | ATCAACAAGTTTGTACAAAAAAGCAGG | Forward primer for amplifying *ftsN*-vector fragment for pJL019 |
| 72 | ACTAGTAGTTAATTTCTCCTCTTTAATG | Reverse primer for amplifying vector fragment for pJL019 and pJL109 |
| 127 | GAGGAGAAATTAACTACTAGTATGGCACAACGAGATTATGTACGC | Forward primer for cloning *ftsN* into pJL098 and pJL119 |
| 128 | GACCCTTAGCGGCCGCTTAACCCCCGGCGGCGAGC | Forward primer for cloning *ftsN* into pJL098 and pJL119 |
| 11 | GGAGAAATTAACTACTAGTATGGTGAGCAAAGGCGAAGAAG | Forward primer for cloning *mNeonGreen* into pJL019 |
| 12 | GCTTTTTTGTACAAACTTGTTGATTTTATACAGTTCATCCATGCCCATC | Reverse primer for cloning *mNeonGreen* into pJL019 |
| 148 | GGTGGCGGTGGTAGCGCACCTTCGCGGCGAAAAAAG | Forward primer for amplifying *ftsN*$^{12\text{-}13}$-vector fragment |
| 149 | GCTACCACCGCCACCCGGTTGGCTGCGGCGTACA | Reverse primer for amplifying *ftsN*$^{12\text{-}13}$-vector fragment |
| 150 | CCAACCGGGTGGCGGTGGTAGCGTGAGCAAAGGCGAAGAAGATAAC | Forward primer for cloning *mNeonGreen* into pJL107 |

| | | |
|---|---|---|
| 151 | CGAAGGTGCGCTACCACCGCCACCTTTATACAGTTC ATCCATGCCCATC | Reverse primer for cloning *mNeonGreen* into pJL107 |
| 141 | GCTACCACCGCCACCATTTCGTTGCTTTTTCCGTGA GGTG | Forward primer for amplifying $ftsN^{28\text{-}29}$-vector fragment |
| 142 | GGTGGCGGTGGTAGCCTGCCTGCGGTTTCTCCCG | Reverse primer for amplifying $ftsN^{28\text{-}29}$-vector fragment |
| 143 | GCAACGAAATGGTGGCGGTGGTAGCGTGAGCAAAG GCGAAGAAGATAAC | Forward primer for cloning *mNeonGreen* into pJL103 |
| 144 | GCAGGCAGGCTACCACCGCCACCTTTATACAGTTCA TCCATGCCCATC | Reverse primer for cloning *mNeonGreen* into pJL103 |
| 152 | GGTGGCGGTGGTAGCGAGTCCGAGACGCTGCAAA G | Forward primer for amplifying $ftsN^{60\text{-}61}$-vector fragment |
| 153 | GCTACCACCGCCACCTTCTTTCTTGTGATGCGTAAT GAAGTAC | Reverse primer for amplifying $ftsN^{60\text{-}61}$-vector fragment |
| 154 | CAAGAAAGAAGGTGGCGGTGGTAGCGTGAGCAAAG GCGAAGAAGATAAC | Forward primer for cloning *mNeonGreen* into pJL108 |
| 155 | CGGACTCGCTACCACCGCCACCTTTATACAGTTCAT CCATGCCCATC | Reverse primer for cloning *mNeonGreen* into pJL108 |
| 156 | GGTGGCGGTGGTAGCGTGACCGGAAACGGACTACC | Forward primer for amplifying $ftsN^{69\text{-}70}$-vector fragment |
| 157 | GCTACCACCGCCACCTTTCTGGCTTTGCAGCGTCTC G | Reverse primer for amplifying $ftsN^{69\text{-}70}$-vector fragment |
| 158 | CCAGAAAGGTGGCGGTGGTAGCGTGAGCAAAGGC GAAGAAGATAAC | Forward primer for cloning *mNeonGreen* into pJL109 |
| 159 | GGTCACGCTACCACCGCCACCTTTATACAGTTCATC CATGCCCATC | Reverse primer for cloning *mNeonGreen* into pJL109 |
| 160 | GGTGGCGGTGGTAGCCTGACACCAGAACAACGTCA GC | Forward primer for amplifying $ftsN^{113\text{-}114}$-vector fragment |
| 161 | GCTACCACCGCCACCTTGCTCCGGCGTTTTCACTTC AC | Reverse primer for amplifying $ftsN^{113\text{-}114}$-vector fragment |
| 162 | CGGAGCAAGGTGGCGGTGGTAGCGTGAGCAAAGG CGAAGAAGATAAC | Forward primer for cloning *mNeonGreen* into pJL110 |

| | | |
|---|---|---|
| 163 | GTGTCAGGCTACCACCGCCACCTTTATACAGTTCATCCATGCCCATC | Reverse primer for cloning *mNeonGreen* into pJL110 |
| 164 | GGTGGCGGTGGTAGCATGCAGGCTGATATGCGCCAG | Forward primer for amplifying *ftsN*$^{124\text{-}125}$-vector fragment |
| 165 | GCTACCACCGCCACCTTGTTCAAGAAGCTGACGTTGTTCTG | Reverse primer for amplifying *ftsN*$^{124\text{-}125}$-vector fragment |
| 166 | GAACAAGGTGGCGGTGGTAGCGTGAGCAAAGGCGAAGAAGATAAC | Forward primer for cloning *mNeonGreen* into pJL111 |
| 167 | CCTGCATGCTACCACCGCCACCTTTATACAGTTCATCCATGCCCATC | Reverse primer for cloning *mNeonGreen* into pJL111 |
| 129 | GCTACCACCGCCACCTTGCTGACGCTGTTCCGGC | Forward primer for amplifying *ftsN*$^{151\text{-}152}$-vector fragment |
| 130 | GGTGGCGGTGGTAGCACGCTACAGCGCCAACGTC | Reverse primer for amplifying *ftsN*$^{151\text{-}152}$-vector fragment |
| 131 | GCGTCAGCAAGGTGGCGGTGGTAGCGTGAGCAAAGGCGAAGAAGATAAC | Forward primer for cloning *mNeonGreen* into pJL100 |
| 132 | GTAGCGTGCTACCACCGCCACCTTTATACAGTTCATCCATGCCCATC | Reverse primer for cloning *mNeonGreen* into pJL100 |
| 133 | GCTACCACCGCCACCCTGCTGCTGCCAGCTTTGTTC | Forward primer for amplifying *ftsN*$^{182\text{-}183}$-vector fragment |
| 134 | GGTGGCGGTGGTAGCACGCGTACGTCGCAAGCCG | Reverse primer for amplifying *ftsN*$^{182\text{-}183}$-vector fragment |
| 135 | CAGCAGCAGGGTGGCGGTGGTAGCGTGAGCAAAGGCGAAGAAGATAAC | Forward primer for cloning *mNeonGreen* into pJL101 |
| 136 | CGTACGCGTGCTACCACCGCCACCTTTATACAGTTCATCCATGCCCATC | Reverse primer for cloning *mNeonGreen* into pJL101 |
| 137 | GCTACCACCGCCACCTTGCAGCAGATCCTGGTACGG | Forward primer for amplifying *ftsN*$^{212\text{-}213}$-vector fragment |
| 138 | GGTGGCGGTGGTAGCACTCCTGCGCACACGACTGC | Reverse primer for amplifying *ftsN*$^{212\text{-}213}$-vector fragment |
| 139 | CTGCAAGGTGGCGGTGGTAGCGTGAGCAAAGGCGAAGAAGATAAC | Forward primer for cloning *mNeonGreen* into pJL102 |

| | | |
|---|---|---|
| 140 | CGCAGGAGTGCTACCACCGCCACCTTTATACAGTTCATCCATGCCCATC | Reverse primer for cloning *mNeonGreen* into pJL102 |
| 39 | AAAAGCAGGCTCCGCGGCCGCCCCCTTCACCAAGTGAGCAAAGGCGAAGAAGATAAC | Forward primer for cloning *mNeonGreen* into pJL028 |
| 40 | GGCTGCAGGTCGACCCTTAGCGGCCGCTTATTTATACAGTTCATCCATGCCCATC | Reverse primer for cloning *mNeonGreen* into pJL028 |
| 126 | GCGGCCGCTAAGGGTCG | Forward primer for amplifying vector fragment for pJL119 |
| 152 | GGTGGCGGTGGTAGCGAGTCCGAGACGCTGCAAAG | Forward primer for amplifying *ftsN*$^{60-61}$-vector fragment |
| 153 | GCTACCACCGCCACCTTCTTTCTTGTGATGCGTAATGAAGTAC | Reverse primer for amplifying *ftsN*$^{60-61}$-vector fragment |
| 177 | CAAGAAAGAAGGTGGCGGTGGTAGCGGATCCGAAATCGGTACTGGC | Forward primer for cloning *Halo* into pJL132 |
| 178 | CTCGGACTCGCTACCACCGCCACCACCGGAAATCTCCAGAGTAGAC | Reverse primer for cloning *Halo* into pJL132 |
| 60 | CCAGAAGAACGCGCTCGCGCCATTAAAGAGCTG | Forward primer for site-specific mutagenesis into WYAA |
| 61 | CAGCTCTTTAATGGCGCGAGCGCGTTCTTCTGG | Reverse primer for site-specific mutagenesis into WYAA |

**Table S4. Summary of different FtsN fusions**

| Construction | Insertion site | Complementation | Doubling time (hr) | Localization[c] |
|---|---|---|---|---|
| MG1655[a] | N.A.[b] | N.A.[b] | 1.7 ± 0.1 | Midcell |
| mNeG-FtsN | N-terminus | Yes | 1.5 ± 0.1 | Midcell |
| P12-mNeG-A13 | Within $FtsN^{Cyto}$ | Yes | 1.8 ± 0.1 | Midcell, weak |
| N28-mNeG-L29 | | Partially | 2.5 ± 0.3 | Midcell, very weak |
| E60-mNeG-E61 | Between $FtsN^{TM}$ and $FtsN^{E}$ | Yes | 1.5 ± 0.1 | Midcell |
| K69-mNeG-V70 | | Yes | 1.5 ± 0.1 | Midcell |
| Q113-mNeG-L114 | Between $FtsN^{E}$ and $FtsN^{SPOR}$ | Yes | 2.0 ± 0.1 | Midcell & Cell pole |
| Q124-mNeG-M125 | | Yes | 1.6 ± 0.1 | Midcell & Cell pole |
| Q151-mNeG-T152 | | Yes | 1.6 ± 0.1 | Midcell & Cell pole |
| Q182-mNeG-T183 | | Yes | 1.5 ± 0.1 | Midcell & Cell pole |
| Q212-mNeG-T213 | | Yes | 1.6 ± 0.2 | Midcell & Cell pole |
| FtsN- mNeG | C-terminus | Yes | 1.4 ± 0.1 | Midcell & Cell pole |

[a] This is MG1655 wild-type cell. Localization data was from the immunostaining fluorescence images.
[b] N.A. not applicable.
[c] Cells were grown in M9-glucose minimum medium.

**Table S5. FtsN-ring and FtsZ-ring dimension measurements**

| Strains | Deconvolved ring width FWHM (nm)[b] | Deconvolved ring thickness FWHM (nm)[b] | *n*[c] |
|---|---|---|---|
| mEos3.2-FtsN | 86 ± 3 | 51 ± 4 | 115 |
| FtsZ-mEos3.2 [a] | 84 ± 2 | 47 ± 2 | 126 |

[a] FtsZ-mEos3.2 data was from a previous work[8].
[b] The FWHM was deconvolved as previously described[30], which allows comparison of dimensions obtained with different spatial resolutions.
[c] *n* is the number of cells used in each measurement.

**Table S6. Comparison of the moving speed across different divisome proteins**

| Proteins[b] | Imaging Modality | $P_1\_V_1$ (%)[c] | $V_1$ (nm $s^{-1}$)[c] | $V_2$ (nm $s^{-1}$)[c] |
|---|---|---|---|---|
| FtsN | TIRF | 100 | 8.7 ± 0.2 | N.A.[a] |
| | SMT | 100 | 9.5 ± 0.2 | N.A.[a] |
| FtsW | SMT | 63.6 ± 7.6 | 9.4 ± 0.3 | 37.8 ± 6.1 |
| FtsI | SMT | 53.9 ± 19.9 | 9.8 ± 1.1 | 31.2 ± 5.6 |
| FtsZ | TIRF | 0 | N.A.[a] | 28.0 ± 1.2 |

[a] N.A. not applicable

[b] FtsN data is from this study, where TIRF data is the combination of TIRF and TIRF-SIM imaging of the mNeG-FtsN fusion (Strain 4564, Table S1), SMT data is from SMT imaging of the FtsN-Halo$^{SW}$ fusion (Strain 5234, Table S1). FtsW data is from SMT imaging of a FtsW-RFP fusion in a previous study[16]. FtsI data is from SMT imaging of an RFP-FtsI fusion from a previous study[16]. FtsZ data is from TIRF imaging of an FtsZ-GFP fusion in a previous study[12].

[c] Speeds of FtsN ($V_1$) and FtsZ ($V_2$) from the TIRF data were calculated as the average of the absolute speeds. Errors are the *s.e.m.* with $n > 200$. Percentage ($P_1\_V_1$), speed ($V_1$) of the slow-moving population and speed ($V_2$) of the fast-moving population from the SMT data of FtsN, FtsW, and FtsI are obtained from one-population (FtsN) or two-population (FtsW, FtsI) free-float fitting of 1000 CDF curves bootstrapped from three experiments. Errors are the standard deviations of the fitted parameters.

**Table S7. FtsN dynamics in cells with different FtsZ treadmilling speeds**

| FtsZ mutants | FtsZ speed[a] (nm $s^{-1}$) ($N_Z$) | *P*_moving[b] (%) ($N_{all}$) | FtsN speed[b] (nm $s^{-1}$) | *T*_moving[b] (s) ($N_m$) | *T*_stationary[c] (s) ($N_s$) |
|---|---|---|---|---|---|
| WT | 28.0 ± 1.2 (182) | 44.0 ± 1.3 (161) | 9.6 ± 0.4 | 16.4 ± 1.1 (71) | 32.9 ± 3.8 (90) |
| E238A | 23.8 ± 2.9 (37) | 43.1 ± 1.7 (149) | 10.5 ± 0.4 | 15.6 ± 1.6 (64) | 20.8 ± 1.8 (85) |
| E250A | 17.4 ± 1.6 (41) | 43.9 ± 1.6 (285) | 9.3 ± 0.3 | 19.3 ± 1.2 (125) | 32.1 ± 2.5 (160) |
| D269A | 14.3 ± 1.3 (32) | 41.8 ± 1.5 (219) | 9.4 ± 0.3 | 17.9 ± 1.1 (92) | 34.3 ± 3.0 (127) |
| G105S | 9.7 ± 1.0 (35) | 42 .5 ± 1.8 (189) | 10.0 ± 0.4 | 14.4 ± 0.8 (80) | 37.8 ± 4.4 (109) |

[a] FtsZ treadmilling speeds were calculated as the average of the absolute speeds from a previous work[12]. Errors are the *s.e.m.* $N_z$ is the number of FtsZ kymograph segments.
[b] Percentage of moving segments (*P*_moving), average moving speed, and average dwell time (*T*_moving) of FtsN molecules. Errors are the standard deviation from 1000 bootstrap samples pooled from three independent experiments. $N_{all}$ is the number of total track segments. $N_m$ is the number of segments corresponding to a directionally moving molecule.
[c] Average dwell time (*T*_stationary) of FtsN molecules in the stationary state. Errors are the standard deviation from 1000 bootstrap samples pooled from three independent experiments. $N_s$ is the number of segments corresponding to a stationary molecule.

**Table S8. Dynamics of FtsN's cytoplasmic domain mutants**

| FtsN mutant | *P*_moving[a] (%) ($N_{all}$) | FtsN speed[a] (nm $s^{-1}$) | *T*_moving[a] (s) ($N_m$) | *T*_stationary[b] (s) ($N_s$) |
|---|---|---|---|---|
| $FtsN^{D5N}$ | 43.1 ± 2.4 (202) | 9.7 ± 0.4 | 11.4 ± 0.6 (87) | 17.7 ± 1.8 (115) |
| $FtsN^{\Delta Cyto\text{-}TM}$ | 40.5 ± 1.6 (234) | 9.2 ± 0.3 | 10.5 ± 0.6 (94) | 27.7 ± 2.5 (140) |

[a] Percentage of segment number (*P*_moving), average speed, and average dwell time (*T*_moving) of FtsN mutant molecules spent in directional moving state. Errors are the standard deviation from 1000 bootstrap samples pooled from three independent experiments. $N_{all}$ is the number of total track segments. $N_m$ is the number of segments corresponding to a directionally moving molecule.
[b] Average dwell time (*T*_stationary) of FtsN mutant molecules in the stationary state. Errors are the standard deviation from 1000 bootstrap samples pooled from three independent experiments. $N_s$ is the number of segments corresponding to a stationary molecule.

**Table S9. FtsN$^{Cyto\text{-}TM}$ dynamics in cells with different FtsZ treadmilling speeds**

| FtsZ mutants | FtsZ speed[a] (nm $s^{-1}$) ($N_Z$) | *P*_moving[b] (%) ($N_{all}$) | FtsN$^{Cyto\text{-}TM}$ speed[b] (nm $s^{-1}$) | *T*_moving[b] (s) ($N_m$) | *T*_stationary[c] (s) ($N_s$) |
|---|---|---|---|---|---|
| WT | 28.0 ± 1.2 (182) | 62.5 ± 1.9 (130) | 29.1 ± 1.7 | 7.5 ± 0.4 (81) | 18.4 ± 1.6 (49) |
| E238A | 23.8 ± 2.9 (37) | 59.5 ± 3.1 (108) | 26.4 ± 1.5 | 7.7 ± 0.4 (64) | 15.6 ± 1.3 (44) |
| E250A | 17.4 ± 1.6 (41) | 54.8 ± 3.0 (96) | 21.7 ± 1.2 | 7.1 ± 0.4 (53) | 12.8 ± 0.7 (43) |
| D269A | 14.3 ± 1.3 (32) | 45.1 ± 3.1 (104) | 15.8 ± 0.7 | 7.3 ± 0.5 (47) | 19.5 ± 2.0 (57) |
| G105S | 9.7 ± 1.0 (35) | 37.5 ± 2.0 (197) | 10.9 ± 0.5 | 8.5 ± 0.5 (74) | 12.5 ± 0.9 (123) |

[a] FtsZ treadmilling speeds were calculated as the average of the absolute speeds from previous work. Errors are the *s.e.m.* $N_z$ is the number of FtsZ kymograph segments.
[b] Percentage of segment number (*P*_moving), average speed, and average dwell time (*T*_moving) of FtsN$^{Cyto\text{-}TM}$ molecules spent in directional moving state. Errors are the standard deviation from 1000 bootstrap samples pooled from three independent experiments. $N_{all}$ is the number of total track segments. $N_m$ is the number of segments corresponding to a directionally moving molecule.
[c] Average dwell time (*T*_stationary) of FtsN$^{Cyto\text{-}TM}$ molecules spent in stationary state. Errors are the standard deviation from 1000 bootstrap samples pooled from three independent experiments. $N_s$ is the number of segments corresponding to a stationary molecule.

**Table S10. FtsN dynamics under different sPG synthesis conditions**

| Genotype | Drug or medium | $P$_moving[a] (%) ($N_{all}$) | FtsN speed[a] (nm $s^{-1}$) | $T$_moving[a] (s) ($N_m$) | $T$_stationary[b] (s) ($N_s$) |
|---|---|---|---|---|---|
| BW25113 | M9-glucose | 44.1 ± 2.2 (161) | 9.6 ± 0.4 | 16.4 ± 1.1 (71) | 32.9 ± 3.8 (90) |
| BW25113, *ftsW*$^{I302C}$ | M9-glucose | 43.7 ± 1.3 (155) | 9.6 ± 0.5 | 13.8 ± 1.0 (68) | 26.3 ± 1.5 (87) |
| | MTSES, M9-glucose | 19.6 ± 1.4 (143) | 9.5 ± 0.5 | 12.7 ± 0.9 (28) | 24.1 ± 2.9 (115) |
| MG1655 | M9-glucose | 44.9 ± 1.6 (571) | 9.4 ± 0.2 | 14.5 ± 0.7 (256) | 27.3 ± 1.3 (315) |
| | Aztreonam, M9-glucose | 10.3 ± 0.9 (760) | 7.9 ± 0.5 | 19.2 ± 1.1 (79) | 42.5 ± 1.2 (681) |
| | Fosfomycin, M9-glucose | 9.9 ± 1.8 (374) | 9.7 ± 0.5 | 15.2 ± 0.9 (37) | 40.9 ± 1.8 (337) |
| | 1XPBS (4% PFA fixed) | 5.2 ± 0.9 (856) | 9.1 ± 0.6 | 15.4 ± 1.0 (44) | 73.7 ± 2.3 (812) |
| MG1655, *ftsI*$^{R167S}$ | M9-glucose | 45.8 ± 2.0 (375) | 9.4 ± 0.3 | 16.0 ± 0.7 (173) | 18.5 ± 0.7 (202) |
| | EZRDM | 48.9 ± 1.0 (154) | 12.3 ± 0.5 | 13.8 ± 0.5 (75) | 14.4 ± 0.8 (79) |
| | EZRDM + UppS | 49.2 ± 1.7 (140) | 13.7 ± 0.5 | 13.2 ± 0.8 (69) | 12.8 ± 0.5 (71) |

[a] Percentage of segment number ($P$_moving), average speed, and average dwell time ($T$_moving) of FtsN molecules spent in directional moving state. Errors are the standard deviation from 1000 bootstrap samples pooled from three independent experiments. $N_{all}$ is the number of total track segments. $N_m$ is the number of segments corresponding to a directionally moving molecule.

[b] Average dwell time ($T$_stationary) of FtsN molecules spent in stationary state. Errors are the standard deviation from 1000 bootstrap samples pooled from three independent experiments. $N_s$ is the number of segments corresponding to a stationary molecule.

**Table S11. The *p*-values of the two-sample Kolmogorov-Smirnov (K-S) test for FtsN and FtsW's directional movement**

| Growth condition | *p* | | |
|---|---|---|---|
| | Speed (*V*) | Moving dwell time (*T*_moving) | Processive running length (PL) |
| EZRDM | 0.214[a] | 0.074[a] | 0.369[a] |
| EZRDM + UppS | 0.052[a] | 0.052[a] | 0.108[a] |

[a] The *p*-values indicate the distributions are not significant different from each other ($p > 0.05$).

**Table S12. Dynamics of FtsN mutants in the superfission strain**

| Genotype | Plasmid | *P*_moving[b] (%) ($N_{all}$) | $P_1$_$V_1$[c] (%) | $V_1$[b, c] (nm $s^{-1}$) | $V_2$[c] (nm $s^{-1}$) | *T*_moving[b] (s) ($N_m$) | *T*_stationary[d] (s) ($N_s$) |
|---|---|---|---|---|---|---|---|
| TB28, *ΔftsN*, *ftsB*[E56A] | $FtsN^{WT}$ | 44.0 ± 1.8 (285) | 100 | 9.3 ± 0.3 | N.A.[a] | 14.8 ± 0.8 (125) | 31.2 ± 2.0 (160) |
| | $FtsN^{WYAA}$ | 11.0 ± 2.1 (252) | 100 | 13.6 ± 1.3 | N.A.[a] | 10.9 ± 0.7 (28) | 21.4 ± 1.4 (224) |
| | $FtsN^{E}$ | 62.9 ± 1.2 (202) | 30 | 8.4 ± 1.9 | 28.8 ± 6.3 | 9.1 ± 0.5 (127) | 14.1 ± 0.9 (75) |

All the experiments were performed with cells grown in M9-glucose medium.

[a] N.A. not applicable

[b] Percentage of segment number (*P*_moving), average speed ($V_1$), and average dwell time (*T*_moving) of FtsN or FtsN mutant molecules spent in directional moving state. Errors are the standard deviation from 1000 bootstrap samples pooled from three independent experiments. $N_{all}$ is the number of total track segments. $N_m$ is the number of segments corresponding to a directionally moving molecule.

[c] Percentage ($P_1$_$V_1$), speed ($V_1$) of the slow-moving population and speed ($V_2$) of the fast-moving population of $FtsN^{E}$ molecules obtained from two-population free-float fitting of CDF curves bootstrapped 1000 times from three independent experiments. Errors are the standard deviations of the fitted parameters.

[d] Average dwell time (*T*_stationary) of FtsN or FtsN mutant molecules spent in stationary state. Errors are the standard deviation from 1000 bootstrap samples pooled from three independent experiments. $N_s$ is the number of segments corresponding to a stationary molecule.
